# *EagleImp*: Fast and Accurate Genome-wide Phasing and Imputation in a Single Tool

**DOI:** 10.1101/2022.01.11.475810

**Authors:** Lars Wienbrandt, David Ellinghaus

## Abstract

**Background:** Reference-based phasing and genotype imputation algorithms have been developed with sublinear theoretical runtime behaviour, but runtimes are still high in practice when large genome-wide reference datasets are used.

**Methods:** We developed *EagleImp*, a software with algorithmic and technical improvements and new features for accurate and accelerated phasing and imputation in a single tool.

**Results:** We compared accuracy and runtime of *EagleImp* with *Eagle2, PBWT* and prominent imputation servers using whole-genome sequencing data from the 1000 Genomes Project, the Haplotype Reference Consortium and simulated data with more than 1 million reference genomes. *EagleImp* is 2 to 10 times faster (depending on the single or multiprocessor configuration selected) than *Eagle2/PBWT*, with the same or better phasing and imputation quality in all tested scenarios. For common variants investigated in typical GWAS studies, *EagleImp* provides same or higher imputation accuracy than the *Sanger Imputation Service, Michigan Imputation Server* and the newly developed *TOPMed Imputation Server*, despite larger (not publicly available) reference panels. It has many new features, including automated chromosome splitting and memory management at runtime to avoid job aborts, fast reading and writing of large files, and various user-configurable algorithm and output options.

**Conclusions:** Due to the technical optimisations, *EagleImp* can perform fast and accurate reference-based phasing and imputation for future very large reference panels with more than 1 million genomes. *EagleImp* is freely available for download from https://github.com/ikmb/eagleimp.

## Introduction

Genotype phasing and imputation has become a standard tool in *genome-wide association studies (GWAS)*, and the accuracy of phasing and imputation generally increases with the number of haplotypes from a reference panel of sequenced genomes [1], with the algorithmic complexity (and thus the runtime) of the imputation process depending on the number of unique haplotypes in each genomic segment of the target samples (GWAS input dataset) and the total number of these segments in the reference panel. State-of-the-art reference-based phasing and imputation algorithms such as *Eagle2* [2], *PBWT* [3] and *minimac4* [4, 5, 6] have been efficiently developed with runtimes better than linear scaling over the size of the reference panel. For example, Das et al. [5] showed that increasing the reference panel size from 1,092 (1000 Genomes Project Phase 1, 27 million variants) to 33,000 individuals (*Haplotype Reference Consortium (HRC)*, 40 million variants) (>40-fold increase in the number of reference genotypes) only increases the phasing and imputation runtime by a factor of 10 (5.3h vs. 51.3h) to impute 100 GWAS samples in one single-threaded process. However, this would still result in a runtime of weeks for a new reference panel with more than one million reference samples. The *UK Biobank (UKB)* has recently made *whole-genome sequencing (WGS)* data from 200,000 individuals available [7], and the newly established *European ‘1+Million Genomes’ Initiative (1+MG)* [8] project is underway, generating WGS data of over one million genomes. The 1+MG project will lead to a further increase in the number of reference genotypes by more than >30-fold compared to the HRC panel benchmarked by Das et al. [5]. Currently, the best solution to perform phasing and imputation for large datasets is to perform parallel processing on large multi-core systems or *high-performance computing (HPC)* clusters with hundreds of CPU-cores, to distribute the computational load as much as possible. Therefore, algorithmic and technical improvements together with existing implementations are needed to ensure that phasing and imputation remain feasible for reference panels with more than one million samples.

To allow phasing and imputation for very large reference panels, while ensuring at least the same phasing and imputation accuracy, we developed *EagleImp*, a software tool for accelerated phasing and imputation. *EagleImp* introduces algorithmic and implementation improvements to the established tools *Eagle2* [2] (phasing) and *PBWT* [3] (imputation) in a single convenient application. By making changes to the algorithm, parameters, and implementation, we were able to speed up the classical 2-step imputation process with *Eagle2* and subsequent *PBWT* by more than a factor of ten for single chromosome analysis and (depending on the parallelisation strategy) at least more than a factor of two for the entire human genome while maintaining or even improving phasing and imputation quality.

*EagleImp* also provides many new convenient features via simple command line parameters, such as a continuous progress report to a file, user pre-selection of per genotype imputation information (genotypes, allele dosage, genotype dosage and/or probabilities), phasing confidences and usage information of input variants in a separate file, variant IDs from the reference in imputation output, automated chromosome chunking (if main memory requirements are too high), lock file support (to enable two or more processes to share CPU resources), detection and handling of ref/alt swaps and/or strand flips, the ability to skip certain parts of the algorithm (e.g. pre-phasing, reverse phasing or entire phasing or imputation) and more. In addition, *EagleImp* supports imputation of chromosome X and Y with automatic partitioning by pseudo-autosomal regions (*PAR*).

## Background

Since early studies have shown that the accuracy of imputation increases significantly when the genotype data contains information on the haplotype phase of heterozygous variants [9], it is common practice to apply a haplotype phasing algorithm to a target input dataset prior to genotype imputation. Among others, the best known phasing tools are *SHAPEIT2* [10] (phasing without reference) and *Eagle2*, whereby the latter is currently used on all prominent imputation servers, such as the *Sanger Imputation Service (SIS)* [11, 1], *Michigan Imputation Server (MIS)* [12, 13] and the newly developed *TOPMed Imputation Server (TOPMed)* [14, 15]. Prominent imputation tools are *IMPUTE V2* [16], *minimac4* [13] and *PBWT* [3].

In developing *EagleImp*, we focused on improving and merging *Eagle2* and *PBWT* into a single tool. Both tools use the equally named *Position-based Burrows-Wheeler Transform (PBWT)* data structure introduced by Durbin [3] as their basis. Its main advantages are the compact representation of binary data and the ability to quickly look up any binary sequence at any position in the data. The runtime complexity is linear to the length of the query sequence, independent of the size of the database. To create a PBWT, the algorithm determines permutations of the input sequences for each genomic site such that the subsequences ending at that site are sorted when read backwards. In our work, we propose further algorithmic and implementation improvements that allow a more efficient use of the PBWT data structure and thus increase the speed of phasing and imputation, while maintaining at least the same accuracy of phasing and imputation.

### *Improvements in* EagleImp

For algorithmic and computational details of the original phasing in *Eagle2* and imputation in *PBWT*, we refer to our **Supplementary 1** and the original publications by Loh et al. [2] and Durbin [3]. Full details of *EagleImp* improvements summarised below can be found in the **Supplementary 2**.

First, the following points summarise the *EagleImp* improvements to the data structure and further technical improvements: (i) We have developed a new `.qref` format for reference data, which significantly improves the reading time of the reference data (**Supplementary 2.1**). (ii) The PBWT data structure of the *condensed reference* (**Supplementary 1.1**) required for each target sample is now stored in a compressed format (**Supplementary Figure 1** in **Supplementary 2.2**), i.e. a binary format (in permuted form) corresponding to the calculated permutation arrays with an index similar to the *FM-index* used for a *Burrows-Wheeler transformation (BWT)* [17] (**Supplementary Listing 1**), to ensure fast generation, compact storage and fast access to the reference data. (iii) Haplotype probabilities are no longer stored in a log-based format and a non-normalised scaling factor is used for the haplotype path probabilities (**Supplementary 2.2**), which only needs to be updated in case of a predictable loss of precision after several path extension operations. In this way, probability calculations remain precise, especially for heavily needed summations of floating point numbers without an otherwise required approximation (as in *Eagle2*) or back-transformation. (iv) The imputation of missing genotypes during phasing is obsolete since the subsequent imputation step imputes missing genotypes for shared variants (between target and reference) in the same way as variants that only occur in the reference. To implement this, we used a tree structure to calculate set-maximal matches (**Supplementary 2.3**). (v) Unlike the original *PBWT* tool, *EagleImp* uses multiple threads for genotype imputation, including the use of multiple temporary output files to reduce the input/output (IO) file bottleneck (**Figure 1** and **Supplementary 2.3**). (vi) We introduced a conversion of genotypes and haplotypes into a compact representation with integer registers and made extensive use of Boolean and bit masking operations as well as processor directives for bit operations (such as *popcount* for counting the set bits in a register) throughout the application which accelerated the computing time significantly (**Supplementary Figure 1** and **Supplementary Listing 2**).

**Figure 1.**
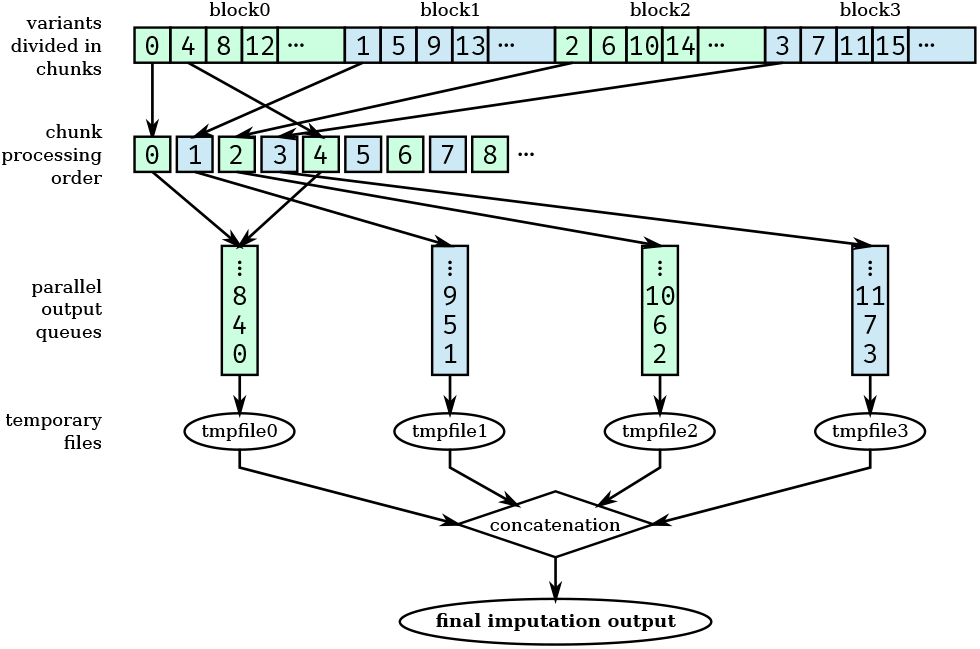
Newly implemented multi-processing scheme for imputation with *EagleImp*. Variants are distributed over blocks with a separate output file. Chunks in a block are processed repetitively iterating over the blocks. Output files are concatenated at the end. The example shows a distribution over four blocks. The numbers in the chunk indicate the order in which they are processed. See **Supplementary 2.3.5** for details.

Second, the following points summarise further algorithmic improvements in *EagleImp*: (i) Due to the properties of the PBWT data structure, sequences that end with equal subsequences at a certain position are located next to each other in the PBWT and can thus be addressed as an *interval*. For the path extension step now only the interval boundaries have to be mapped to the next position to get the corresponding intervals for both possible extensions of the sequence (**Figure 2** and **Supplementary Listing 2**). Since the frequency of a subsequence is equal to the (normalised) size of the corresponding interval, the frequency calculation could thus be accelerated. (ii) We increased the default beam width parameter (number of possible paths) from 50 (fixed value in *Eagle2*) to 128 and the Δ parameter from 20 (fixed value in *Eagle2*) to 24 in favor of further increasing the phasing quality with minimal loss of computing time (**Supplementary 1.1 and 2.2**). (iii) We omitted the *identical-by-descent (IBD)* check performed by *Eagle2* before the phasing process, since we found no loss in phasing quality without the IBD-check implemented. (iv) Pre-phasing is disabled by default (unlike *Eagle2*), as benchmarks showed no improvement (**Supplementary 3.4**). If desired, pre-phasing can be explicitly enabled with a user option in *EagleImp*. Likewise, reverse phasing can be optionally disabled.

**Figure 2.**
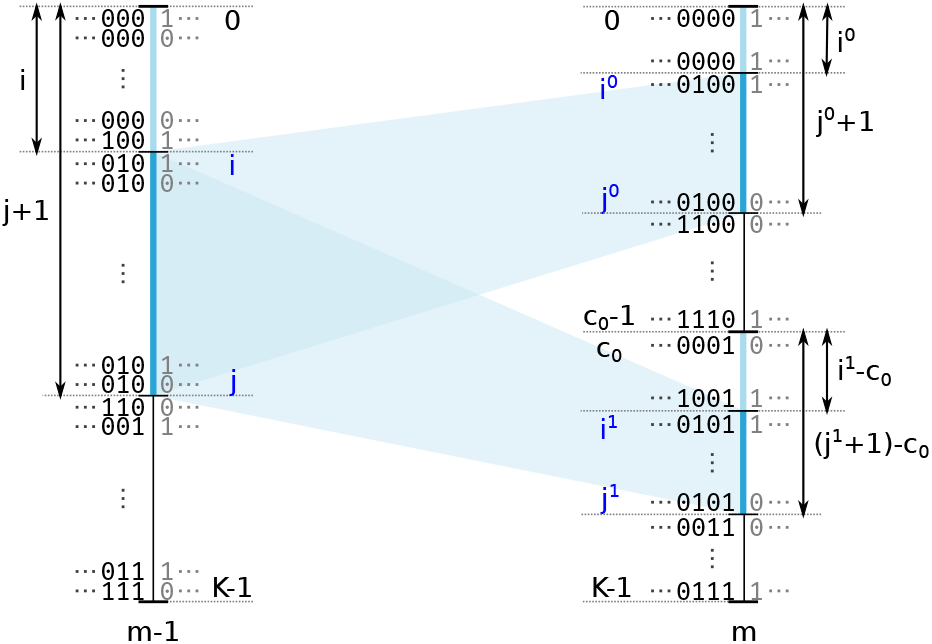
Improved path extension step in *EagleImp*. Illustration of mapping a PBWT interval [*i, j*]_*m*−1_ from position *m* − 1 to *m*, resulting in two intervals [*i*^0^, *j*^0^]_*m*_ and [*i*^1^, *j*^1^]_*m*_. The interval [*i, j*]_*m*−1_ exemplary represents the sequences in the PBWT that end with 010 at position *m*−1. By stepping forward to position *m* these sequences are extended either with 0 or 1 and thus are located in one of the mapped intervals [*i*^0^, *j*^0^]_*m*_ or [*i*^1^, *j*^1^]_*m*_ respectively. It is easy to see, that *i* = *i*_0_ + (*i*_1_ − *c*_0_) and *j* + 1 = (*j*_0_ + 1) + ((*j*_1_ + 1) − *c*_0_) (as *j* is inclusive in the interval) which directly leads to the mapping equations **(6)-(7)** in **Supplementary 2.1**.

Third, we introduced new features and improvements in usability, among other things: (i) *EagleImp* allows imputation of X and Y chromosomes by careful handling of haploid samples. Since *pseudo-autosomal regions (PAR)* are usually encoded with reference to chromosome X in the target data, we often face the problem of diploid data in the PAR regions and haploid data in the non-PAR region of chromosome X target (input) files. Existing tools and imputation servers sometimes crash with an error message if diploid and haploid data is mixed in a single input file. We provide a script together with the *EagleImp* source code that takes care of these regions by automatically splitting the input data with respect to the chromosome X and Y PAR regions before imputation and then merging the imputation results back into one file afterwards. (ii) *EagleImp* allows reference and alternative alleles in the target to be swapped compared to the reference (with the option `--allowRefAltSwap`), e.g. an A/C variant is considered as C/A, and it allows strands to be flipped, e.g. an A/C variant is considered a T/G variant at the same chromosomal position (with the option `--allowStrandFlip`). (iii) *EagleImp* computes the imputation accuracy *r*^2^ (as described in Das et al. [18] and used in *minimac4*) and provides the value together with the allele frequency, the minor allele frequency, the allele count and number as well as the reference panel allele frequency (if available) in the imputation output. An optional *r*^2^ filter can be applied to filter out variants with low imputation quality. (iv) Phasing confidences and information about how the target variants are used for imputation are provided in separate output files. (v) To save disc space for the output files, the user can decide which information is provided along with the imputed (hard called) genotypes, i.e. any combination of allele dosages (*ADS* tag), genotype dosages (*DS* tag), genotype probabilities (*GP* tag) or no information. Variant IDs in imputation output are provided exactly as they appear in the reference. (vi) *EagleImp* automatically activates chromosome chunking if the memory requirement is higher than the available main memory (provided as the runtime parameter `--maxChunkMem`) to eliminate the tedious process of trying out chunk sizes on different input and reference datasets for the user. (vii) For better workload distribution on multi-core computers, we provide a locking mechanism (via a lock file) such that low CPU-load tasks (e.g. reading input files) can run multiple processes at once, while high CPU-load tasks (e.g. the phasing and imputation processes) require multiple-exclusion of CPU resources. We provide an optional launch script that uses this feature for simultaneous processing of multiple input files. (viii) A progress indicator shows the progress in percentage (giving the user a hint how long the analysis will take). Optionally, constantly updated status and info files display summarised information about the running process.

## Data Description

To quantify phasing and imputation quality and runtime improvements of *EagleImp* compared to the original tools *Eagle2* and *PBWT*, first, we conducted quality benchmarks with different parameters on three different sized reference panels (**Table 1**) and 18 target datasets (**Table 2**) including two real-world target GWAS datasets (from [19]) to further compare imputation accuracy of *EagleImp* with the accuracy of current imputation servers (*SIS, MIS* and *TOPMed*). Then, we ran all runtime benchmarks using the *HRC.EUR* target dataset (**Table 2**) and the parameters used for the quality benchmarks. Full descriptions about reference and target datasets from **Tables 1** and **2** and details about the setup for quality and runtime comparisons (in particular the preparation of various multiprocessor configurations) can be found in **Supplementary 3**.

**Table 1.** Three different sized reference panels (A–C) were used for phasing and imputation benchmarks.

|  | Reference | #Variants | #Samples | Ancestry |
| --- | --- | --- | --- | --- |
| (A) | *HRC1.1* | 40.4 million | 27,165 | mixed |
| (B) | *1000G Phase 3* | 84.8 million | 2,504 | mixed |
| (C) | *synthetic HRC1.1* | 40.4 million | 1,086,600 | mixed |

**Table 2.** 18 target datasets in total were used for phasing and imputation benchmarks to evaluate the runtime and impuation accuracy of *EagleImp* compared to the original tools *Eagle2* and *PBWT*. Target datasets (1–6) were sampled from the *HRC1.1* panel (A) and target datasets (7–16) were sampled from the *1000GPhase3* panel (B). For each of the target datasets (1–16), the selected samples were removed from the corresponding reference panels (A) and (B) in the benchmarks to avoid a biased result. The real-world datasets (17–18) (from [19]) did not show an overlap of samples with any of the reference panels.

|  | Target | #Variants | #Samples | Ancestry |
| --- | --- | --- | --- | --- |
| (1) | *HRC.EUR* | 619,872 | 494 | European |
| (2–6) | *HRC.v1–5* | 619,872 | 500 | mixed |
| (7–11) | *1kG.EUR.v1–5* | 647,963 | 50 | European |
| (12–16) | *1kG.v1–5* | 647,963 | 50 | mixed |
| (17) | *COVID.Italy* | 559,519 | 2,113 | Italian |
| (18) | *COVID.Spain* | 549,696 | 1,792 | Spanish |

### Choice of phasing and imputation parameters

The exact input parameters for the program call of *EagleImp, Eagle2* and *PBWT* are listed in **Supplementary 3.3**. For performance comparison, we focused on testing different values of the parameter *K* (to select the *K*-best haplotypes from the reference for phasing). For example, we ran a benchmark on each *HRC*.* dataset (**Table 2** (1–6)) with four different values of *K* for *EagleImp* as well as for *Eagle2:* 10,000 (default setting in *Eagle2*), 16,384, 32,768, and “*max*” (where max means using all available haplotypes for phasing, which is 54,330 minus the number of removed haplotypes (988 for target (1) and 1000 for targets (2–6)) in the case of the reduced *HRC1.1* panel (A)). All runs used one phasing iteration (reflecting the default setting of *Eagle2* if the number of target samples is less than half of the number of reference samples).

### Benchmark system and program versions

The computing system we used consists of two Intel Xeon E5-2667 v4 CPUs, each with 8 cores running at 3.2 GHz, resulting in 32 available system threads. The system is equipped with 256 GB of DDR4 RAM and uses a ZFS file system that combines six HDDs, each with a capacity of 2 TB in a *raidz2* pool (leading to a total capacity of about 7 TB). The operating system is Ubuntu Linux 21.04 with kernel 5.11.0.

*EagleImp* is written in C++ and compiled with GCC v10.3.0. For *Eagle2* and *PBWT* we used the to date most recent builds on Github: *Eagle2 v2.4.1* [20] and *PBWT 3.1-v3.1-7-gf09141f* [21]. Both tools were also compiled with GCC v10.3.0 together with the required *HTSlib v1.12* and *bcftools v1.12*. For runtime benchmarks, we measured the wall-clock runtime by marking the start and end points of the benchmark with the command date and calculated the difference in runtime.

### Metrics for phasing and imputation accuracy

We counted a *phase switch* whenever the current phase at a call site differs the phase at the previous call site, and we counted a *switch error* whenever a phase switch occurs at a calling site in the target but not in the reference when comparing a phased haplotype pair from the target to the original haplotype pair in the reference, or vice versa. The *switch error rate* per sample is then computed by dividing the switch errors by the number of target variants. In our benchmarks, we showed switch error rate for each benchmark set averaged over all samples.

*Genotype errors* are determined in a similar way by comparing the genotypes of an imputed haplotype pair with the corresponding genotypes in the reference panel and counting the number of differences. The number of genotype errors in a sample divided by the number of total variants gives the *genotype error rate*. As with the switch error rate, we calculated the genotype error rate for each benchmark set averaged over all samples.

Another way to determine the imputation quality is the imputation accuracy *r*^2^, which is a valuable means of interpretation with regard to sample size and statistical power in a GWAS study and which can basically be considered independent of minor allele frequency (MAF) (although the precision of the *r*^2^ estimate decreases with low MAF) [18]. *r*^2^ can be estimated for each imputed variant from posterior allele probabilities without knowing the true allele on each chromosome.

## Phasing and Imputation Quality

### *Effect of pre-phasing and reverse phasing on* EagleImp

Contrary to our expectations, pre-phasing in *EagleImp* resulted in a slight loss of quality for all values of *K* for the *HRC1.1* reference panel (an increased switch error and genotype error rate of about 0.06% and 0.1%, respectively, (**Supplementary Tables 1– 2**, **Supplementary Figure 2**) and at the same time increased the runtime (**Supplementary Table 3**), we disabled pre-phasing in *EagleImp* by default. (However, the user can explicitly switch it on again by using the `--doPrePhasing` switch). In contrast, disabling reverse phasing results in a significant loss of quality for all *K* (between 3.2% and 3.7% increased switch error rate and a genotype error rate increased by approximately 1.0% to 1.2% (**Supplementary Tables 1–2**, **Supplementary Figure 2**), and should therefore be retained in *EagleImp*.

### Switch and genotype error rates

For both *EagleImp* and *Eagle2/PBWT*, average (genome-wide) switch error rates and genotype error rates decreased (as expected) with higher values of *K* using the *HRC1.1* reference panel (**Figure 3; Supplementary Tables 4–5**). For *K* = 10,000 the phasing and imputation quality of *EagleImp* is nearly equal to *Eagle2/PBWT* (ranging from an increase of 0.5% and a reduction of 0.2% in the switch error rate (**Figure 3 (a)**), and an increase of 0.5% and a reduction of 0.5% in the genotype error rate **Figure 3 (e)**)). As the value of *K* increases, the *EagleImp* runs performed better compared to the corresponding *Eagle2/PBWT* runs with the same value of *K* (**Figures 3 (b–d)** and **Figures 3 (f–h)**). For example, for *K* = 16,384 the switch error rate is lowered between 0.4% and 0.9%, while for *K* = max, the switch error rate is lowered between 2.2% and 2.4%. For the genotype error rate we measure a reduction from 0% to 0.8% for *K* = 16,384 and from 1.1% to 1.8% for *K* = max. In our *1000G Phase 3* reference benchmark analysis (**Figure 4, Supplementary Tables 6–9**), *EagleImp* performed better than *Eagle2/PBWT* for all 10 target datasets from the *1000G Phase 3* panel.

**Figure 3.**
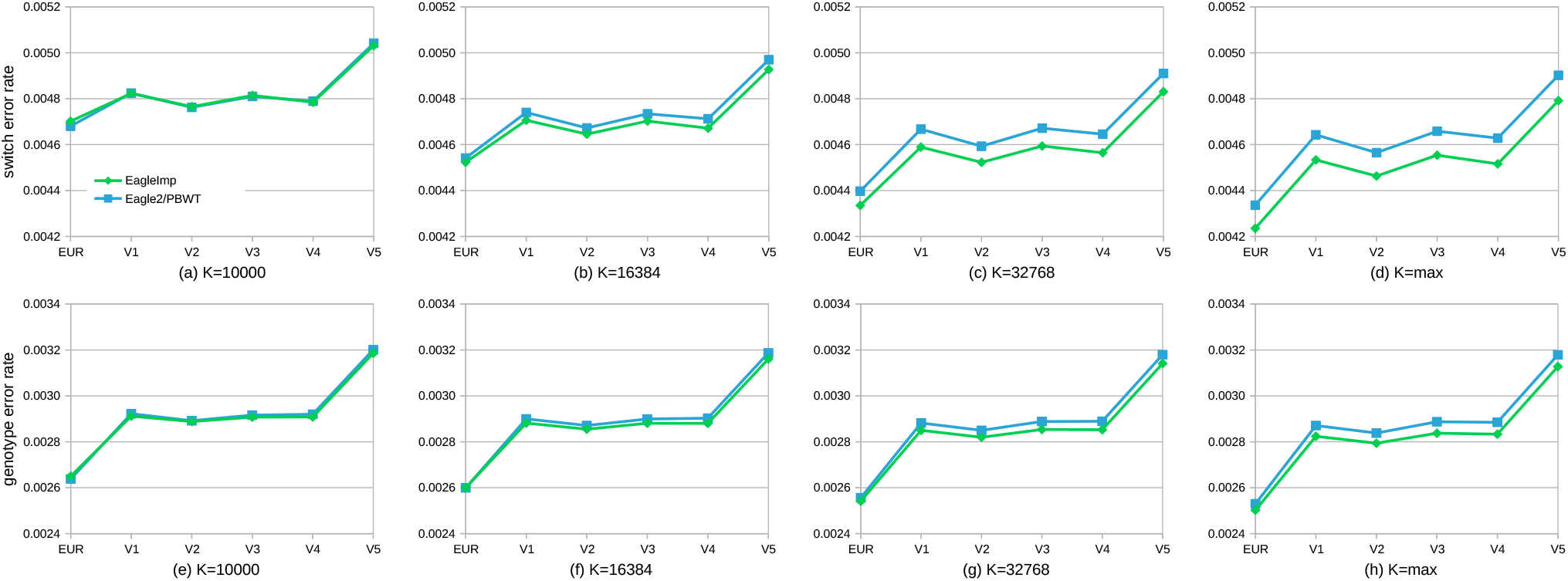
*HRC1.1* reference panel (mostly European ancestry; **Table 1 (A)**) based phasing and imputation performed better for *EagleImp* (green) compared to the original *Eagle2/PBWT* (blue) with increasing *K* in all six test datasets: **(a–d)** Average (genome-wide) switch error rates (after phasing) and **(e–h)** genotype error rates (after imputation) using different values of *K* ∈ {10, 000 (default setting in *Eagle2*); 16, 384; 32, 768; max} for six target datasets named *HRC.EUR* (European ancestry; **Table 2 (1)**) and *HRC.v1–5* (Mixed ancestry; **Table 2 (2–6)**). The parameter *K* selects the K-best haplotypes from the reference for phasing, resulting in better quality but longer runtimes.

**Figure 4.**
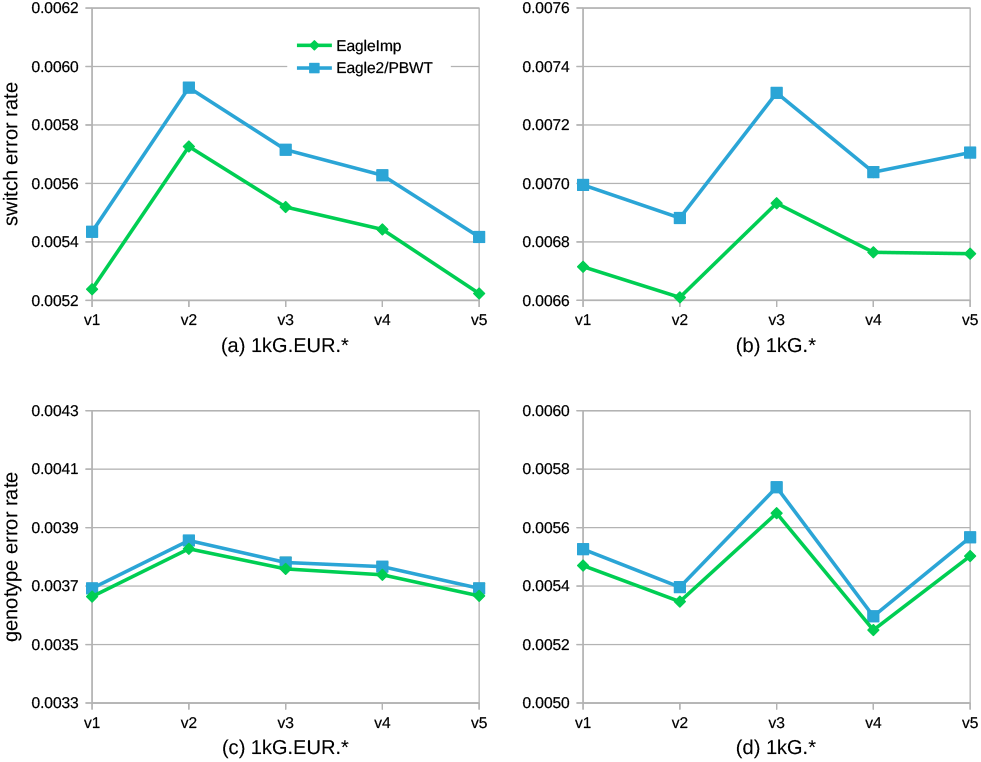
For the *1000G Phase 3* reference panel (23 global populations; **Table 1 (B)**), phasing and imputation with *EagleImp* (green) outperformed the original *Eagle2/PBWT* (blue) in all ten test datasets: **(a–b)** Average genome-wide switch error rates (after phasing) and **(c–d)** genotype error rates (after imputation) for the target datasets named *1kG.EUR.v1–5* (European ancestry; **Table 2 (7–11)**) and *1kG.v1–v5* (different worldwide populations; **Table 2 (12–16)**). Parameter *K* was chosen to be maximum in all test runs (here, *K* = 5, 008), thus including the entire *1000G Phase 3* reference panel.

Our benchmarks with the *HRC1.1* and the *1000G Phase 3* reference panel showed that switch error and genotype error rates were generally lower in the European input target datasets. As expected, due to the smaller reference panel, the *1000G Phase 3* benchmarks reveal higher error rates than in the *HRC1.1* benchmarks. The target datasets with mixed populations showed an increased switch error rate of about 25% while the genotype error rate is increased by around 40% to 50%. When compared to *Eagle2/PBWT, EagleImp* clearly performed better with a reduction of the switch error rate between 4%–5% and the genotype error rate between 0.5%–1%.

### Imputation accuracy r^2^

For the real-world GWAS datasets *COVID.Italy* and *COVID.Spain*, we determined *r*^2^ values stratified by MAF (including all variants above this threshold) using the four different *K* parameters from above (**Supplementary Figures 3** and **4**). The *COVID.Spain* dataset showed a generally slightly better imputation performance than the *COVID.Italy* dataset in all runs, possibly due to a slightly similar genetic background compared to the *HRC1.1* reference panel. Already for *K* = 10,000 (**Supplementary Figure 3a** and **4a**), phasing and imputation with *EagleImp* consistently produced higher *r*^2^ values than the *Eagle2/PBWT* combination across the entire MAF spectrum, and also higher *r*^2^ values than the *SIS*, despite its larger (not freely available) *HRC1.1* reference panel as compared to our *HRC1.1* benchmark panel. With *K* = 10, 000 and a MAF greater than 0.001 (and especially at higher MAF), *EagleImp* also showed a quality advantage in contrast to the *MIS*, which we attribute to our algorithmic changes. Below a MAF of 0.001, the MIS still seems to show its higher imputation quality due to the larger *HRC1.1* reference panel. Especially in this low frequency range, *TOPMed* imputation (as a reference model) shows that a much larger reference panel plays a another key role in increasing imputation accuracy for rare variants, in addition to algorithmic improvements. Interestingly, for *K* = 10, 000 and common variants such as in GWAS studies, *EagleImp* was shown to even achieve a higher quality over *TOPMed* (here for MAF greater than 0.02 for *COVID.Spain* and greater than 0.006 for *COVID.Italy*) probably due to *EagleImp’s* algorithmic improvements, although *EagleImp’s* reference panel is more than three times smaller than that of *TOPMed*, despite the smaller reference panel in both comparisons.

For higher *K* parameters, the graphs for *EagleImp* and *Eagle2/PBWT* showed (as expected) better *r*^2^ values but the distance between both tools remained the same (**Supplementary Figures 3 (b–d), 4 (b–d)**). **Figure 5** depicts the average genome-wide *r*^2^ values for the *COVID.Italy* and *COVID.Spain* datasets as a function of different MAF thresholds (on logarithmic scale), with *K* = max for *Eagle2* and *EagleImp* (which is effectively *K* = 54,330 as we used the public *HRC1.1* panel). Here *EagleImp* achieved at least equivalent or better results compared to the *MIS* across the entire MAF spectrum and showed same quality or a quality gain compared to *TOPMed* for MAF greater than 0.01 (*COVID.Spain*) and 0.004 (*COVID.Italy*).

**Figure 5.**
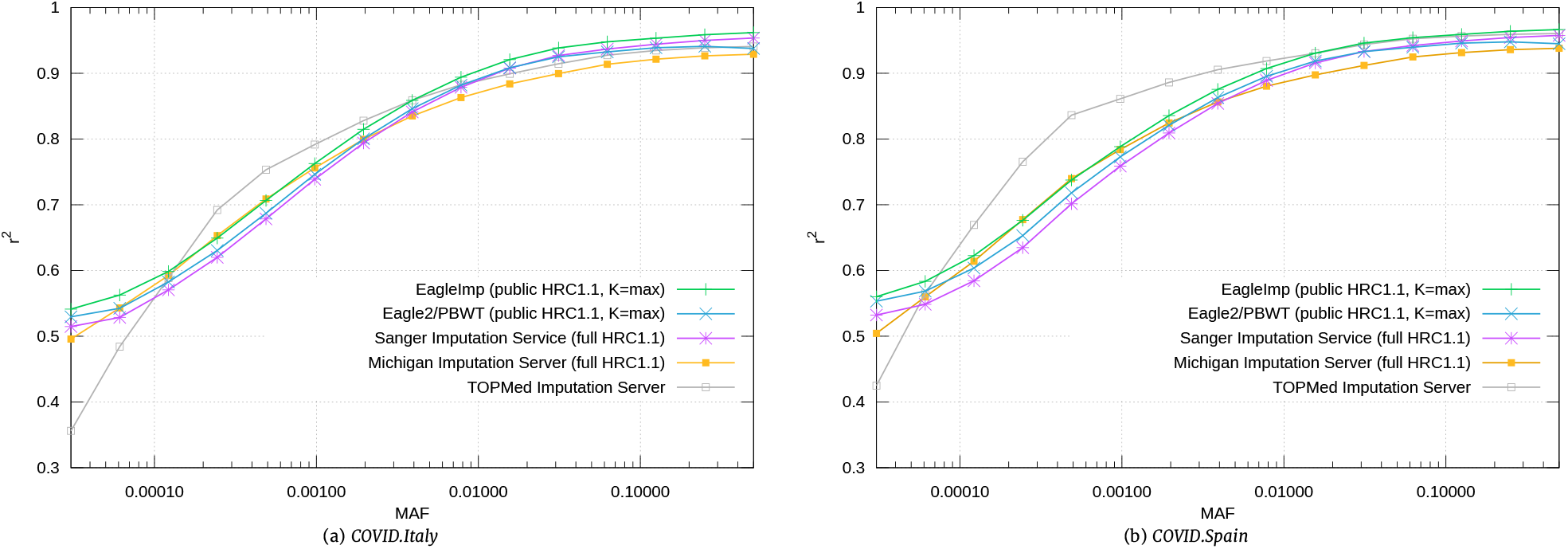
Imputation accuracy *r*^2^ was higher for *EagleImp* (green; *K* = 54, 330 (max)) compared to *Eagle2/PBWT* (blue; *K* = 54, 330 (max)) for two real-world GWAS datasets (a) *COVID.Italy* and (b) *COVID.Spain* (**Table 2 (17–18)**), with further comparison to imputation *r*^2^ from imputation servers *SIS* (purple), *MIS* (orange) and *TOPMed* (grey). *EagleImp* showed a quality gain compared to *TOPMed* for minor allele frequency (MAF) greater than 0.004 (*COVID. Italy*) and 0.01 (*COVID.Spain*) and achieved at least equivalent results compared to *MIS* across the entire MAF spectrum, although *MIS/SIS* and *TOPMed* use their own reference panels, which are approximately 1.2 times (32,470 samples) and 3.5 times (97,256 samples) larger than the publicly available *HRC1.1* panel (27,165 samples; **Table 1 (A)**) used for *EagleImp* and *Eagle2/PBWT*.

## Phasing and Imputation Runtime

For the sake of simplicity, we ran all runtime benchmarks using the *HRC.EUR* target dataset and the parameters used for the *HRC1.1* reference benchmarks above, but with different multiprocessing configurations with up to 32 concurrent system threads for *EagleImp* and *Eagle2/PBWT*. Details of the various multiprocessor configurations tested can be found in **Supplementary 3.7**. In addition, we examined runtimes of individuals chromosomes processed with all 32 system threads (referred to as *1×32* runs) (**Supplementary 3.8**) and single-thread runtimes with multi-processing disabled (**Supplementary 3.9**), since multi-processing strategies can behave differently on different computing systems. We also measured the runtimes of our real-world GWAS datasets *COVID.Italy* and *COVID.Spain* which can be found in **Supplementary 3.10**. Note, that the runtime benchmarks do not include the preparation of the reference files (**Supplementary 3.11**), which is required for *PBWT* and is optional for *EagleImp. (PBWT* requires a `.pbwt` file for each reference file; for *EagleImp* we used our newly developed `.qref` format instead of `.vcf.gz` or `.bcf`.)

### Multi-processing runtimes

We observed that for any value of *K*, the *Eagle2.8×4* configuration performed best for the *Eagle2/PBWT* runs and the *EagleImp.2×4×8* configuration performed best for *EagleImp* (**Supplementary Table 10**), which is why we only compared these two configurations in the following.

For *K* = 10, 000 (**Figure 6 (a)**), *Eagle2.1×32* took 1 hour and 46 minutes: The *Eagle2.8×4* configuration accelerated this by a factor of 2.30 to 46 minutes. In contrast, the *EagleImp.1×32* configuration required 40 minutes, which we could speed up by a factor of 1.85 to 22 minutes in the *EagleImp.2×4×8* configuration. This corresponds to a speedup factor of 2.66 when comparing the *1×32* configurations of *Eagle2/PBWT* and *EagleImp* or a factor of 2.14 when comparing the fastest multi-processing configurations of the two tools. The total runtime increased with higher values of *K* (**Figure 6 (b–d)**), but the speedup factors between both tools only varied slightly. For *K* = max, the *Eagle2/PBWT* runtime was 2 hours and 27 minutes for the *1×32* configuration and 1 hour and 25 minutes for *8×4*. *EagleImp* analysed the same data in 54 minutes (*1×32*) and 35 minutes (*2×4×8*), resulting in speedup factors of 2.71 and 2.41, respectively.

**Figure 6.**
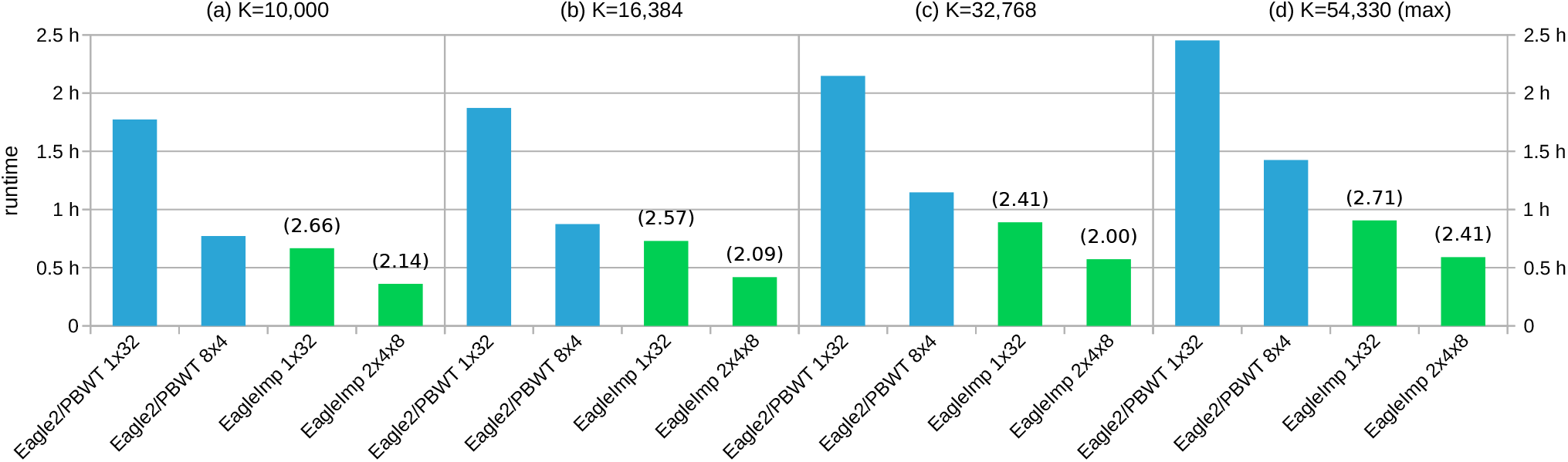
Faster runtimes of *EagleImp* compared to *Eagle2/PBWT* (with at least the same accuracy, **Figure 3**), demonstrated for the processing of 494 GWAS target samples using the *HRC1.1* reference panel (**Table 1(A)**) and different *K* parameters (*K*=54330 denotes the maximum value including all haplotypes from the HRC1.1 panel). The naive multi-processor configuration (*1×32* with 32 phasing threads and each chromosome processed sequentially) and the best individual multiprocessor configuration (i.e. fastest individual configuration determined from 6 different runs of *Eagle2/PBWT* and 8 different runs of *EagleImp*, see **Supplementary Table 10**) were compared. The numbers in brackets give the acceleration factor of *EagleImp* compared to the corresponding *Eagle2/PBWT* run.

When comparing runtimes for individual chromosomes, we additionally measured runtimes for phasing-only, imputation-only and combined runs. For the phasing-only runs, we observed that *EagleImp* is between 2.93 and 4.81 times faster than *Eagle2* for all chromosomes and different values of *K* (**Supplementary Table 11**). The imputation-only runs of *EagleImp* with 32 threads when compared to *PBWT* showed the advantages of the multi-threading capability of *EagleImp* (which *PBWT* does not offer), with a speedup between 7.77 and 12.94 (**Supplementary Table 12**). The combination of phasing and imputation offers the advantage for *EagleImp* that the reference panel does not have to be read twice (as is the case with *Eagle2* and *PBWT*), resulting in a combined speedup between 5.82 to 10.81 for single chromosomes (**Supplementary Table 13**).

### Single-thread runtimes

We measured the runtimes of the *HRC.EUR* dataset from above with the four values of *K* exemplary for chromosome 2 and chromosome 21 (largest and smallest number of input variants) with only one thread for *Eagle2* and *EagleImp* (by using the corresponding `--numThreads` parameters for both tools). We found that *EagleImp* is faster than *Eagle2/PBWT* with measured speedups between 1.59 and 2.48 for phasing only and between 1.40 and 1.51 for imputation only, and a combined speedup between 1.58 and 2.25 (**Supplementary Tables 14–15**).

## Phasing and Imputation using One Million Reference Genomes

To our knowledge, the *TOPMed* imputation server uses the largest reference panel currently available for phasing and imputation, with 97,256 reference samples (*TOPMed r2*) [15]. To explore *EagleImp*’s capabilities in handling a future reference panel with more than one million samples, we created a synthesised panel *synthetic HRC1.1* (1,086,600 samples and 40.4 million variants, **Table 1 (C)**) by multiplying the sample data of the public *HRC1.1* panel (**Table 1 (A)**) 40 times (see section *Data Description*). We measured *EagleImp*’s and *Eagle2/PBWT*’s runtimes for the real-world GWAS dataset *COVID.Italy* (2,113 samples) using *K* = 10, 000 and *K* = 2, 173, 200 (maximum *K* from the *synthetic HRC1.1* panel) and the naive (*Eagle2.1×32, EagleImp.1×32*) and respectively fastest multi-processor configurations (*Eagle2.8×4, EagleImp.2×4×8*) from the previous runtime benchmark (**Supplementary Table 18**).

*EagleImp* outperformed *Eagle2/PBWT* with a speedup factor of at least 2.80 (*K* = 10,000) and 3.50 (*K* = max) (**Figure 7**). *EagleImp* took about 24 hours for *K* = 10, 000 in *2×4×8* configuration, while, in contrast, the original *Eagle2/PBWT* took almost three days (68 hours) in *8×4* configuration respectively. The runs with *K* = max, i.e. *K* = 2,173,200 in this case, resulted in a runtime of 4 days for *EagleImp* in *2×4×8* configuration, while *Eagle2* could not complete the run due to insufficient memory and missing automatic chunking ability (requirement of more than 256GB of RAM) on our benchmark system in *8×4* configuration. In the *1×32* configuration *Eagle2* crashed only for the larger chromosomes 1–4 and 6 with a total runtime (including the crashes) of more than 15 days.

**Figure 7.**
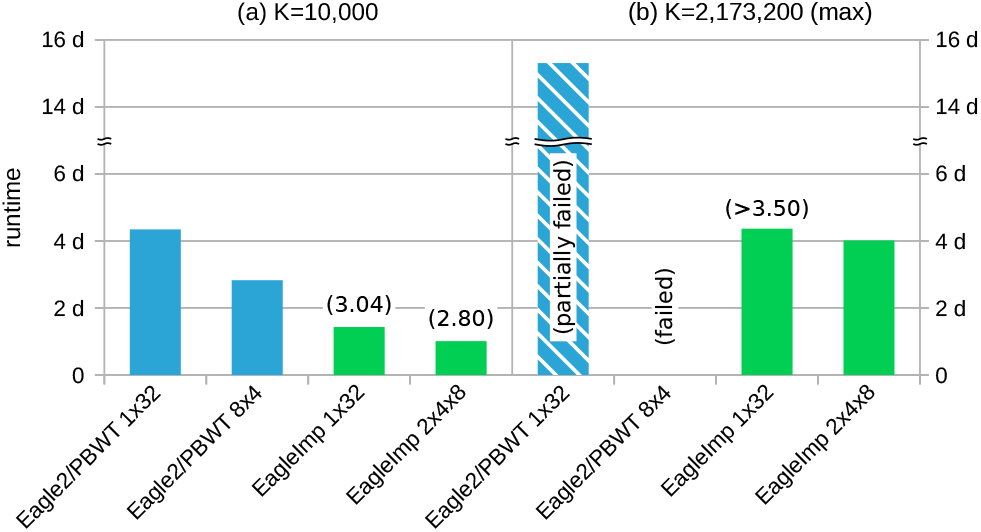
Phasing and imputation runtimes for the *synthetic HRC1.1* reference panel (**Table 1 (C)**) containing >1 million samples (2,173,200 haplotypes) and the real-world *COVID.Italy* dataset (2,113 samples; **Table 2 (17)**). The numbers in brackets give the acceleration factor of *EagleImp* compared to the corresponding *Eagle2/PBWT* run. For *K* = max *Eagle2* could not complete the analysis in *8×4* configuration due to insufficient main memory (marked with “failed”), hence, no runtime or speedup could be measured in this configuration. For the *1×32* configuration phasing with *Eagle2* crashed for chromosomes 1–4 and 6 (“partially failed”) and the stated runtime includes the incomplete runs.

## Discussion

We have introduced *EagleImp*, a fast and accurate tool for phasing and imputation. Due to technical improvements and changes in the data structure, *EagleImp* is 2 to 10 times faster (depending on the multi-processor configuration) than *Eagle2/PBWT*, with same or better phasing and imputation quality in all tested scenarios. For common variants investigated in typical GWAS studies, *EagleImp* also yielded equal or higher imputation accuracy than the imputation servers *MIS*, *SIS* and *TOPMed* that use larger (not freely available) reference panels, which we attribute to our algorithmic improvements. Because of the technical optimisation and the improvement of the stability of the software, *EagleImp* can perform phasing and imputation for upcoming very large reference panels with more than 1 million genomes.

For phasing, we accelerated the search for the *K* best haplotypes, the generation of the condensed reference and the entire phasing process by using an alternative haplotype encoding, Boolean comparison operations and processor directives for bit operations. We improved the accuracy of the phase probability calculation by using a scaled floating point representation instead of a logarithm-based representation, and we improved the frequency lookups in the PBWT data structure by introducing our interval mapping procedure and omitting duplicate calculations of frequencies in the beam search. We investigated the effects of pre-phasing and reverse phasing procedures with the conclusion that pre-phasing is unnecessary. By introducing multi-threading and writing multiple temporary files during imputation, and by using a tree-structure and the same interval mapping technique as for the phasing part to search for set-maximal matches, we were able to speed up the imputation process considerably compared to *PBWT*. Furthermore, the correct treatment of missing genotypes from the input dataset became possible with the tree-structure.

An additional reduction in computing time was made possible by the introduction of the *Qref*-format (producible from standard `.vcf` or `.bcf` files) for fast reference loading, by combining phasing and imputation in a single tool, and by an helper script that introduces multi-processing with several worker processes and a balanced distribution of a complete genome-wide input dataset. We further enhanced the usability of phasing and imputation by adding several convenience features, such as chromosome X and Y handling, imputation *r*^2^ and *MAF* calculation for output files, user selection of desired imputation information (allele dosages, genotype dosages, genotype probabilities), automated optimisation of chromosome partitioning (chunking) and memory management during runtime, preserving variant IDs from reference, *r*^2^ filtering and other things.

Unfortunately, we were not able to sufficiently investigate the quality of the imputation of rare variants, as we did not have the larger reference panels of the *MIS*, *SIS* and the *TOPMed* imputation servers at our disposal. We noticed that in the imputation of common variants, the *MIS* (uses *minimac4* for imputation) fell behind the *SIS* (uses *PBWT* for imputation) in terms of quality, despite the same larger HRC reference panel, but the *MIS* performed better than the *SIS* in the imputation of rare variants. We have not investigated this further. However, since new, even larger reference panels will be available soon, this difference should not matter too much. We already designed *EagleImp* to use more than one million reference genomes (>2 million haplotypes) for reference-based phasing and imputation. In this case, *EagleImp* will be able to show its full strength in terms of runtime and quality compared to other tools, although again an optimisation of the *K* parameter (selection of the *K* best reference genomes where a higher *K* increases the runtime but produces better results) will be required. Imputation in combination with new methods such as study-specific pre-selection of reference samples using deep learning imputation reconstruction methods for reference panels [22] could provide a higher accuracy for rare variants.

As an improvement, in the future we plan to replace the slow *HTSlib* with our own library in order to speed up the writing process since this is still a bottleneck in *EagleImp*. A reduction of the runtime by using *Field Programmable Gate Arrays (FPGAs)* is another possibility to reduce the runtime even further (Wienbrandt et al. [23]). Furthermore, we plan to offer a free web service for *EagleImp*, so that the advantages of *EagleImp* can also be used by the community even without special hardware equipment.

## Methods

We give detailed information about the original concepts of *Eagle2* [2] and *PBWT* [3] in **Supplementary 1** and describe our changes to the algorithm and implementation in **Supplementary 2**, with additional details about the performed benchmarks in **Supplementary 3**.

## Availability of source code and requirements

- Project name: EagleImp
- Project home page: https://github.com/ikmb/eagleimp
- Operating system: Linux
- Programming language: C++ (bash, awk)
- Other requirements: HTSlib, Zlib, BOOST, TBB, Cmake
- License: GNU GPL v3.0

## Availability of supporting data and materials

All quality and runtime measures from our benchmarks are listed in the **Supplementary Material**. The *1000 Genomes Phase 3* reference panel can be downloaded at ftp://ftp.1000genomes.ebi.ac.uk/vol1/ftp/release/20130502/. Data access to the EGAD00001002729 dataset for the HRC1.1 panel is restricted and was granted under request ID 11699. The benchmark datasets *HRC.EUR, HRC.v1–5,1kG.EUR.v1–5* and *1kG.v1–5* are subsets of the previously mentioned reference panels.

## Declarations

## List of abbreviations

BCF: Binary Variant Call Format
BWT: Burrows-Wheeler Transform
HRC: Human Reference Consortium
PBWT: Position-based Burrows-Wheeler Transform
VCF: Variant Call Format

## Ethical Approval

All participants provided written informed consent, and the study was approved by the ethics boards of the participating institutions in agreement with the Declaration of Helsinki principles.

## Author’s Contributions

**Lars Wienbrandt** Application software development; hardware infrastructure; benchmark setup, execution, analysis, interpretation; manuscript writing
**David Ellinghaus** Data acquisition; benchmark analysis, interpretation; manuscript writing; project supervision

## Acknowledgements

The authors would like to thank the COVID-19 GWAS Group for the use of the COVID-19 GWAS data for the benchmark.

