## Supplementary Material for "*EagleImp*: Fast and Accurate Genome-wide Phasing and Imputation in a Single Tool"

January 11, 2022

### 1 Original Core Algorithms of *Eagle2* and *PBWT*

Here, we briefly describe the core algorithms of the original tools *Eagle2* and *PBWT*. For details, we refer to the original publications [1] and [2].

#### 1.1 *Eagle2* Phasing Core Algorithm

The phasing core of *Eagle2* consists of some preliminary steps before the actual phasing, which are performed for each target sample independently.

##### 1.1.1 $K$ best haplotypes

At first, a subset of the reference of the best-fitting haplotypes is determined. These haplotypes have the lowest *genotype-to-haplotype* distance to the sample. (This distance is simply calculated by counting the number of inconsistent haplotype alleles to the corresponding target genotype.) The number is defined by the input parameter $K$. A smaller $K$ reduces the number of haplotypes selected from reference panel and thus reduces phasing quality in favor of a faster runtime. *Eagle2* used $K = 10,000$ by default as a good compromise between quality and speed. This step is only omitted if $K$ is greater than the total number of available reference haplotypes. In this case simply all haplotypes are taken.

##### 1.1.2 Condensed reference and PBWT data structure

The set of the $K$ best haplotypes is reduced to *call sites* afterwards. These are the sites where a phase call is required (basically heterozygous target sites). The information stored in the *condensed reference* is the allele of each haplotype at each call site and the information if a segment between call sites is *consistent* with the target or not, i.e. if the segment between two call sites (implicitly containing only homozygous sites) matches the corresponding segment in the haplotype according to the genotype-to-haplotype distance.

After this step the original *Eagle2* method introduces a so-called *IBD-check (identical by descent)*. This check is intended to remove a certain bias introduced by reference sequences that have a high similarity to the target (caused by a probable relatedness). Segments in the condensed reference are simply marked inconsistent if runs of consistent segments are too long (the default parameters are 10 and 20 segments depending on two different types of IBD).

Based on the condensed reference a *PBWT* data structure is created in order to perform quick sequence lookups from each call site, which is required by the following phasing process. For details on the PBWT, we refer to Durbin [2].

##### 1.1.3 Phasing

The original phasing method in *Eagle2* is categorised into three sub parts: Fast *pre-phasing*, *fine phasing* and *reverse phasing*. All three parts are handled subsequently by performing a *beam search* with different parameters. The parameters for the pre-phasing part are runtime optimised. It is intended to constraint phases with very high probability. This information is later used in the fine phasing part to reduce its runtime. Here, the phase decisions are made by calculating the *keep-phase* probability $P^{\text{keep}}$ for each call site (see "Beam search and probability model" below). The reverse phasing part is intended to fix the phasing decisions from the fine phasing and is performed only in the last iteration (in the case several phasing iterations are performed). In this part, a phase correction is made if and only if the re-caclulated probability for a phase keep or switch is higher than the previously calculated one in the fine phasing.

##### 1.1.4 Beam search and probability model

A *beam* consists of several pairs of haplotype paths at each position in the condensed reference. The probability of such a pair (further referred to as *diplotype*) forms the basis of the keep-phase probability at a call site: If the sum of the probabilities of all diplotypes in the beam at a call site that keep the current phase is higher than for those introducing a phase switch, the current phase is kept, otherwise switched.

The probability model for a diplotype is the following:

$$P\left(h^{\text{mat}}_{1\ldots m}, h^{\text{pat}}_{1\ldots m}|g_{1\ldots m}\right) \approx \underbrace{P\left(g_{1\ldots m}|h^{\text{mat}}_{1\ldots m}, h^{\text{pat}}_{1\ldots m}\right)}_{\varepsilon^{n_{\text{err}}}} P\left(h^{\text{mat}}_{1\ldots m}\right) P\left(h^{\text{pat}}_{1\ldots m}\right) \tag{1}$$

$n_{\text{err}}$ is the number of consistency errors between the pair of haplotype paths to the target genotypes, e.g. if both paths show a common allele at a site (homozygous) while the genotype is heterozygous at that site, this counts as an error. $\varepsilon$ is a fixed error probability set to 0.003 per default.

The haplotype path probability is assumed to follow:

$$P\left(h_{1\ldots m}\right) \approx \sum_{x=m-H}^{m-1} P\left(h_{1\ldots x}\right) f\left(h_{x+1\ldots m}\right) P\left(\text{rec } m|x\right) \tag{2}$$

The parameter $H$ indicates the size of a *history*, i.e. how many sites should be looked back at in order to calculate the actual probability of the current site $m$. This defaults to 30 for the pre-phasing step and is set to 100 in the fine and reverse phasing steps. $P\left(\text{rec } m|x\right)$ is the recombination probability between the two sites $m$ and $x$. It is computed as a function dependent on the genetic distance between the sites, the effective population size and the number of references. The function $f(h_{x\ldots m})$ returns the frequency of the given sequence $h_{x\ldots m}$ in all reference sequences from sites $x$ to $m$. Since this function is called very frequently a fast execution is crucial. The PBWT data structure allows this information to be accessed in approximately constant time while a naive implementation would be linear in the number of references.

The keep-phase probability can now be quantified as the sum of all probabilities of diplotypes in the beam that keep the phase divided by all diplotype probabilities:

$$P^{\text{keep}} = \frac{\sum\left\{P_d\left(h^{\text{mat}}_{1\ldots m}, h^{\text{pat}}_{1\ldots m}|g_{1\ldots m}\right)\middle| d \text{ keeps phase}\right\}}{\sum P_d\left(h^{\text{mat}}_{1\ldots m}, h^{\text{pat}}_{1\ldots m}|g_{1\ldots m}\right)} \tag{3}$$

The *switch-phase* probability is quantified as its counterpart $P^{\text{switch}} = 1 - P^{\text{keep}}$. The *phasing confidence* for a complete sample, which is reported at the end of the phasing part, is the average of all phase confidences for each call site, i.e. either the keep-phase probability if the phase is kept, or the switch-phase probability otherwise.

The beam search algorithm sweeps over the call sites while the beam is updated continuously. *Eagle2* uses a *HapHedge* data structure for this purpose which is basically a tree with the leaf nodes representing active beam paths. Diplotypes that have a probability below $\varepsilon$ times the current maximum diplotype probability are removed. Furthermore, the least probable diplotypes are removed until the number of diplotypes is less than or equal to the beam width parameter, which defaults to 50 in general (30 for pre-phasing).

A parameter $\Delta$ lets the beam advance to site $m+\Delta$ before doing the phase call for site $m$. This ensures that only stable diplotypes that survive until $m+\Delta$ in the beam are taken for calculating the phase probability. This defaults to 20 (10 for pre-phasing).

#### 1.2 *PBWT* Imputation Core Algorithm

##### 1.2.1 Set-maximal matches

Preliminary to imputation, the core algorithm requires the reference to be in PBWT structure format. Together with the target data a reference panel in PBWT format reduced to common variant positions is created. Then, for each sample haplotype all *set-maximal matches* are calculated. A *match* is defined by an interval $\left[m^{\text{start}}, m^{\text{end}}\right)$ (inclusive start and exclusive end positions of the match in the reduced panel) where the target haplotype matches a set of reference haplotypes exactly. A match is set-maximal to a sequence in the reference, if the match is locally maximal, i.e. there is no longer match including this one, and there is also no other sequence in the reference with a longer match that includes this region. Note, that set-maximal matches may (and should) overlap. It is worth to mention that the original PBWT tool sets missing target haplotypes to the reference allele without further notice in the default mode. Missing data may be imputed explicitly as an extra step before the final imputation only, and in this step only the phased target data is used for the imputation instead of the reference panel. In *EagleImp* we do not follow this approach (see **Section 2.3.2**).

##### 1.2.2 Dosage calculation

For each variant allele that needs to be imputed, the imputation dosage is calculated as follows. For each sequence (indexed with $i$) from a set-maximal match that covers this variant, a score $s_i$ is computed that depends on the position of the variant in the match and the length of the match:

$$s_i = \left(m - m_i^{\text{start}}\right)\left(m_i^{\text{end}} - m\right) \tag{4}$$

whereby $m$ is the position of the last common variant in the reduced reference panel and $m_i^{\text{start}}$ and $m_i^{\text{end}}$ the start and end positions of the corresponding set-maximal match. Note, that this equation produces a zero score for all variants in the first segment in a set-maximal match (between the first and the second common site in the reduced panel). In *EagleImp* we add 1 in the first factor to generate a positive score instead (see main paper).

The allele dosage is simply calculated as:

$$d = \frac{\sum\{s_i | i \text{ with allele at } p \text{ is alt.}\}}{\sum s_i} \tag{5}$$

In the case that variants are not covered by set-maximal matches, PBWT simply fills these gaps with the most frequent allele and sets the dosage to the reference panel allele frequency.

### 2 *EagleImp* Implementation

*EagleImp* combines the basic algorithmic concepts of *Eagle2* [1] and *PBWT* [2] described above. In the following, we describe our main variations in the algorithm and implementation, divided into a general part with methods addressing the complete application, the part regarding only the phasing step, and finally the imputation part.

#### 2.1 General Improvements

##### 2.1.1 *Qref* reference format

Reading large reference input panels is slow if the panel is provided in VCF-format (i.e. `.vcf.gz` or `.bcf`). On the one hand, VCF is a very flexible format for storing genetic data, but on the other hand, it becomes cumbersome when only certain parts of the provided information needs to be parsed. Thus, several imputation tools require their own formats, e.g. `.m3vcf` for *minimac4* [3] or `.pbwt` for the *PBWT* tool [4]. This circumvents the processing power required for parsing VCF data and the data is ideally already in the desired format for the tool. To be compatible with other tools, we still support the reference panel in VCF-format, but as reference panels are reused several times for different target datasets, we targeted this runtime bottleneck with our own *Quick Reference Format (Qref).*

The Qref-format is basically a consecutive arrangement of the required metadata of the input file followed by the haplotype data. The metadata are firstly numerical values, such as the number of haplotypes and the number of variants, followed by all variant positions, then all variant IDs, all reference allele IDs, all alternative allele IDs, allele frequencies and flags that mark multi-allelic variants (see below). For chromosome X, flags that mark haploid reference samples are also stored in a Qref. Choosing this format, the reading process can directly assign the data to already prepared memory areas without further restructuring and parsing.

The haplotype information, which is the largest part by far, is stored binary encoded in a variant-major format. To save memory, one line of haplotype information is converted to a binary sequence which is run-length encoded afterwards. We use a run-length encoding that uses 8 bit patterns (7 bits for the run-length and 1 bit for the data that is repeated). In most cases, the run-length encoding leads to a significant reduction in size, but in those cases where an encoded line is not smaller than the original, we simply store the not-encoded sequence.

When creating a QRef-file multi-allelic variants are converted to bi-allelic in advance by splitting. However, these variants are flagged such that a user may distinguish them from originally bi-allelic variants and is able to exclude them later with the `--excludeMultiAllRef` switch. A `.qref` file can easily be created from a VCF-file by applying the `--makeQref` switch in *EagleImp*.

##### 2.1.2 PBWT operations

The PBWT data structure is required by the phasing process and the imputation process. For phasing, the PBWT is generated from the condensed reference for each target (once for the pre-phasing and fine phasing steps in forward direction and once in reverse direction for the reverse phasing step). For imputation a single PBWT is required for the complete reference which is then accessed by all targets.

In its basic form the PBWT requires a permutation array $a$ for each position which results in huge memory requirements. Encouraged by Durbin [2] we create and store PBWT data structures required for the phasing and imputation part in a compact format with an index similar to the *FM-index* for a *Burrows-Wheeler transform (BWT)* [5]. **Supplementary Listing 1** shows the pseudo-code of the creation of a PBWT from a reference in our format. The index simply indicates for each position the number entries that are 0 after each block of 32 entries. Note, that the original PBWT algorithm determines the *count0*-value (i.e. the total number of 0-entries for the current position) during creation of the PBWT. The disadvantage is that two arrays (one of length *count0*, the other of length $N-count0$) have to be merged after each position. In *EagleImp* we can either determine these values beforehand on-the-fly during pre-processing (e.g. while reading the reference data) or we compute it very quickly taking benefit from our Boolean data representation and the processor directive *popcount*. Either way, we can omit array merging as we know the start position of the second array beforehand, which in turn saves computation time.

Also note, that we prepare the input data in transposed (i.e. variant-major) format, as the outer loop of the algorithm iterates over the sites and for each site fast access on the haplotype data of several sequences is required.

Supplementary Listing 1: PBWT creation from $N$ Boolean sequences of length $M$.

```
input haps[M][N] # haplotype data
input cnt0[M] # number of 0-entries for each site
output pbwt[M][N] # PBWT permuted haplotype data
output idx[M][N/32] # index: 0-entries after each 32 entries
define a[N] # current permutation array
define b[N] # next permutation array

# Initialise PBWT with all zeros
pbwt[0..M-1][0..N-1] = 0

# Initialisation of initial permutation with the identity
for n in [0..N-1] do
  a[n] = n
done

# loop over all sites
for m in [0..M-1] do
  p0 = 0
  p1 = cnt0[m]
  for n in [0..M-1] do
    if haps[m][a[n]] == 0 then
      # current entry in permutation order is 0
      b[p0] = a[n]
      p0 = p0 + 1
    else
      # current entry in permutation order is 1
      b[p1] = a[n]
      pbwt[m][n] = 1
      p1 = p1 + 1
    fi
    # store index value every 32 entries
    if n%32 == 31 then
      idx[m][n/32] = p0
```

```
    fi
  done
  swap a,b
done
```

For faster access we store the PBWT data and the index in an interlaced format. In particular, each 32 bits of PBWT data are stored in a 32bit-integer followed by the corresponding index value also stored in a 32bit-integer.

The basic operation required on the PBWT is to identify where a certain sequence is positioned in the permutation order of a specific site $m$ if we know the sequence's positional index $i$ at the previous site $m-1$. This operation should ideally be in constant time complexity as it is required for a fast frequency lookup of certain sequences in the phasing step (see below). Basically, this is a lookup of the relative permutation of index $i$ at site $m-1$. However, since storing these permutation arrays would irrationally increase the memory requirements for the PBWT, we have to decode this value from the compact form created from **Supplementary Listing 1**. Clearly, the relative permutation of $i$ must be the exact number of 0-entries until $i$ at site $m-1$ if the corresponding data bit is 0. This is easily to be calculated from the PBWT index value at the field corresponding to $i$ minus the number of all 0-entries subsequent to $i$ until the index value was stored (as it is stored every 32 entries only). Analogue, if the data bit is 1, the relative permutation must be the exact number of 1-entries until $i$ at site $m-1$ plus the number of all 0-entries (as 1-entries are stored after all 0-entries), which can be calculated likewise. **Supplementary Listing 2** shows a simplified pseudo-code to determine the relative permutation index of an index $i$ from site $m-1$ to $m$.

Supplementary Listing 2: Lookup of relative permutation index in PBWT of an index $i$ from site $m-1$ to $m$.

```
input i # index to decode the relative permutation index
input pbwt[N] # PBWT data at site m-1
input idx[N/32] # PBWT index at site m-1
input cnt0 # number of 0-entries for site m-1
output relp # relative permutation index

# 32bit subsequence of PBWT containing i
# encoded in a 32bit integer
s = (int) pbwt[i/32..i/32+31]

# the position of interest (i) is marked with 1
mask = 1 << (i%32)
# extract data
bit = s AND mask
# all preliminary positions excluding i are 1
mask = mask - 1
# subtract the remaining zeros
s = s OR mask
fcnt0 = idx[i/32] - popcount(NOT s)

if bit == 0 then
  # return the number of 0-entries until i
  relp = fcnt0
else
  # return the number of 1-entries until i
  # plus all 0-entries
  fcnt1 = i - fcnt0
  relp = cnt0 + fcnt1
fi
```

Note, that this lookup procedure is clearly in constant runtime complexity and uses fast Boolean operations and the processor directive *popcount*.

Based on this, we implement a fast lookup procedure for frequencies of certain sequences when iterating over the sites of a PBWT. The frequency lookups are required by the core algorithms in phasing and imputation, and for an efficient lookup we deviate from the procedures described by Durbin [2]. Since a main feature of the PBWT structure is that at each position $m$ in the PBWT the prefixes are sorted backwards, the number of occurrences $l$ of a sequence $h_{x\ldots m}$ in the PBWT at $m$ can be determined from an interval $[i_x, j_x]_m$ with $l = j_x - i_x + 1$ (the size of the interval) and $i_x$ and $j_x$ pointing to indices in the actual permutation at $m$ in the PBWT such that the corresponding reference sequences equal $h_{x...m}$ at positions $x$ to $m$ (and sequences before $i_x$ and after $j_x$ do not).

When advancing from a position $m-1$ to $m$ the calculation of an interval $[i_x, j_x]_m$ from the predecessor $[i_x, j_x]_{m-1}$ is simple, if above condition applies: As the corresponding sequences at $m-1$ match $h_{x...m-1}$ and the extension $h_m$ can only be either 0 or 1, mapping $[i_x, j_x]_{m-1}$ must result in two intervals $[i_x^0, j_x^0]_m$ and $[i_x^1, j_x^1]_m$ depending on the value at $h_m$. Thus, we only need to know where the interval borders map to (which we directly get from the permutation array) and we can calculate the borders of the other interval directly. Say w.l.o.g. $i_x$ is mapped to $i_x^0$, and $j_x$ is mapped to $j_x^0$ (i.e. both interval borders have a 0-allele at the new position) and let $c_0$ be the number of zero-bits at $m$. Then the alternative interval $[i_x^1, j_x^1]_m$ can be calculated as follows:

$$i_x^1 = i_x - i_x^0 + c_0 \quad (6)$$

$$j_x^1 = j_x - j_x^0 + c_0 - 1 \quad (7)$$

The calculation is analog if $i_x$ or $j_x$ were mapped to $i_x^1$ or $j_x^1$. **Figure 2** in the main paper illustrates this relation.

#### 2.2 Phasing Core Improvements

As explained above, we keep the main concept of the *Eagle2* core algorithm for phasing. In particular, phasing is divided into phasing preliminaries and the three steps fast pre-phasing, fine phasing and reverse phasing. We also take over the probability model and the concept of the *beam search* (see supplement or [1]). (According to our benchmarks (see **Section 3.4**), we disable the pre-phasing step per default, with a user option to explicitly re-enable it again.)

##### 2.2.1 *K* best haplotypes and the condensed reference

Changes in the phasing procedure include the extensive use of Boolean operations in the selection of the *K* best haplotypes and the creation of the condensed reference in the phasing preliminaries.

The reduction to common variants in target and reference is already done while loading the input data. As the preliminaries contain procedures that work either sample-wise or variant-wise, we store the reduced reference data twice in sample-major and in variant-major format.

***K* best haplotypes** For the selection of the *K* best haplotypes, we require the reference data in sample-major format. In general, each allele in a haplotype is encoded as a bit (`0` = reference, `1` = alternative). An allele sequence forming a haplotype is encoded in a vector of 64bit-integers, such that one integer can store up to 64 alleles. (Multi-allelic variants in the target sample are excluded from phasing while the user may allow multi-allelic reference variants, although these would be split into bi-allelic variants then.) Genotypes are encoded in two bits per genotype, one bit indicating if the genotype is homozygous reference (`is0`) and the other indicating if the genotype is homozygous alternative (`is2`). If both bits are not set, the genotype is heterozygous, while both bits set indicate a missing genotype. A genotype sequence is stored in a vector of 64 bit integers (as for haplotypes) with the difference that each two subsequent integers form a pair where the first integer stores 64 bit of `is0` information, while the second stores `is2` information. With this encoding, choosing the *K* best haplotypes can be performed very quickly using Boolean operations for comparing genotypes and haplotypes. The vector of inconsistencies (`inc`) from a haplotype (`hap`) to a genotype (encoded in `is0` and `is2`) is calculated as follows:

$$\texttt{inc} = (\texttt{is0}\,\text{AND}\,\texttt{hap})\,\text{OR}\,(\texttt{is2}\,\text{AND NOT}\,\texttt{hap}) \quad (8)$$

The *genotype-to-haplotype* distance is then determined by using the processor directive *popcount* that returns the number of set bits in an integer, which is equal to the number of inconsistencies in this case. The *K* sequences with the smallest distance finally form the *K* best haplotypes. Note, that **Supplementary Equation (8)** produces an inconsistency if the genotype is missing. Anyway, this is not a problem since this inconsistency is generated for every reference sequence and thus does not influence the order of best haplotypes.

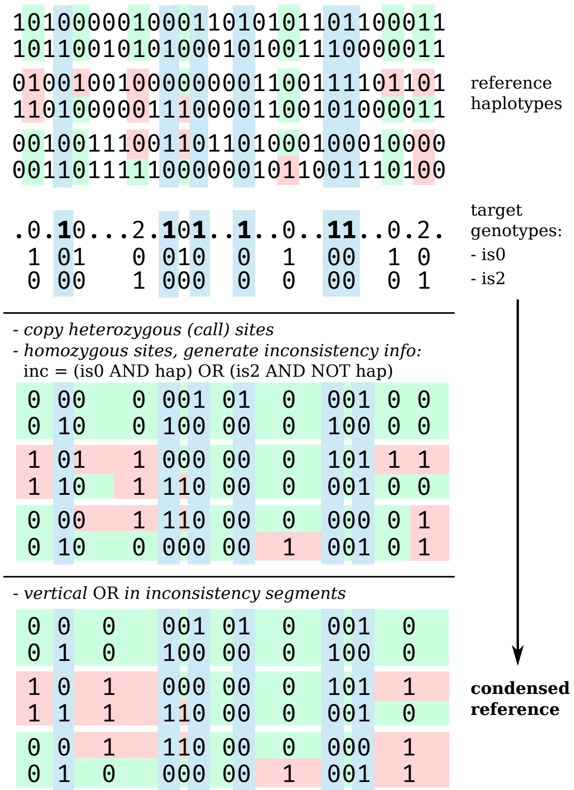

Supplementary Figure 1: In *EagleImp*, we introduced a bit representation of the data (target and reference) packed into 64bit-integers, allowing efficient use of Boolean operations to create the condensed reference. The illustration shows an example of creating a condensed reference for a target sample prior to phasing. Common call sites (heterozygous genotypes) are marked in blue. Haplotypes consistent with the target are marked in green in the reference as well as consistent segments in the condensed reference. Inconsistent haplotypes and the resulting inconsistent segments are marked in red.

**Condensed reference** For creating the condensed reference, the variant-major reference format comes in handy. For each target split site the haplotype information from the reference is copied (selective from the $K$-best haplotypes or simply all information if $K$ is greater or equal to the number of reference haplotypes). For the inconsistency segment between two split sites a complete inconsistency vector of the current target to all references is generated. This is generally faster then selecting the $K$ best haplotypes first as all Boolean operations can be performed on full 64bit blocks instead of selecting $K$ references for each site in the segment first. The inconsistency vector is simply calculated by a consecutive bitwise OR operation with each inconsistency vector for each site in the segment. The inconsistencies for a single site are easily determined similar to **Supplementary Equation (8)** with the difference that the current target genotype is a constant now for the current site. Thus, the inconsistencies are equal to the reference haplotypes themselves if the genotype is homozygous reference or their negation if the genotype is homozygous alternative. For missing genotypes we consider all references to be consistent, so no operation is required at all. The same would apply for heterozygous genotypes, but this case can be excluded as heterozygous target sites have to be selected as split sites before. Only after reaching the next split site (or the end respectively) the $K$ best haplotypes are selected from the complete inconsistency vector and copied to the condensed reference. An example of a condensed reference and its creation is depicted in **Supplementary Figure 1**.

##### 2.2.2 Beam search

A major difference to *Eagle2* phasing applies for the *beam search* where *Eagle2* applies a *HapHedge* data structure, which we omit in favor of a simple list data structure to manage active haplotype and diplotype paths. This is mainly achieved by our interval mapping technique (see above) and the fact that all active paths have to be extended in each iteration (over the sites) anyway, which is why a data structure with a fast forward iteration is favourable. As haplotype path extensions are easily to be foreseen (extensions may be either 0 or 1), it is easy to keep the list sorted by the path's history of haplotypes. And as *beam paths* (or *diplotype paths*) only manage pointers to the underlying haplotype paths, the required merging and pruning steps in the *beam search* can be easily implemented using the sorted haplotype path list and fast Boolean operations for matching paths.

Furthermore, due to the sort order of the path history, equal PBWT intervals are organised next to each other. This characteristic can be used to prevent unnecessary multiple PBWT lookups in the extensions that require interval mappings.

##### 2.2.3 Haplotype Path Probabilities in the Beam Search

The core equation for calculating the phase probability is **Supplementary Equation (2)** that computes a haplotype path probability. The most performance critical step here is the frequency lookup $f(h_{x+1...m})$ since it is called up to $H$ times for each path (whereby $H$ is the history parameter that defaults to $H = 100$ and $x$ increasing from $m-H$ to $m-1$). $f$ returns the (relative) number of occurrences of the haplotype path $h_{x+1...m}$ in the reference from positions $x+1$ to $m$. So, ideally, the process to lookup the frequency has to be in constant time complexity, which we achieve by our interval mapping technique described in **Section 2.1.2** above.

To compute the path probability in **Supplementary Equation (2)**, we need the history of recent path probabilities $P(h_{1...x})$ and the PBWT intervals $[i_{x+1}, j_{x+1}]_m$ to $[i_m, j_m]_m$ that lead to the frequencies $f(h_{x+1...m})$. The history is processed from the newest (most recent) to the oldest site. The recombination probabilities $P(\text{rec } m|x)$ are computed only once for each site according to Loh et al. [1]. Note, that the condensed reference alternately stores call site and segment inconsistency information. To advance from one call site to the next thus implies two PBWT interval mappings, whereby from the first mapping (to the inconsistency information) only the 0-interval (containing only consistent sequences) will be taken in the second mapping.

We observed that a lot of haplotype paths in the beam equal each other in the most recent sites. Thus, we ordered the path extensions such that paths that match each other at the last sites are located next to each other. This way, we can save computing time by directly copying the interval mappings of the previous paths for those positions that are equal, which can be determined quickly by comparing the actual path only to its predecessor and only to the first non-matching position.

**Floating point format** Haplotype path probabilities may become extremely small, especially within a multiplicative combination to a beam path probability (see **Supplementary Equations (1)** and **(2)**). The original *Eagle2* tool solves this problem by transforming the probability values into a logarithm-based representation. In this representation multiplications are solved by addition and additions should be solved by back-transformation first. *Eagle2* circumvents the problem of back-transformation by performing an approximation for the addition of two probabilities in log-based format. We find this solution problematic, as in the computation of the haplotype path probability (**Supplementary Equation (2)**) a summation over up to $H$ terms is required, whereby in-turn each term was computed the same way before. This leads to a strong error propagation which we addressed in *EagleImp* by keeping the probabilities in the non-transformed format but introducing a separate exponential scaling factor, such that the real probability $P$ is defined by the scaling factor $\alpha$ and the internal probability representation $P_{\text{int}}$:

$$P(h_{1...m}) = 2^{\alpha}\ P_{\text{int}}(h1\ldots\text{m}) \tag{9}$$

To save computation time, we keep the scaling factor non-normalised and do an update only after several path extension operations in the case the probability drops below a certain limit. In addition, we keep the scaling factor equal throughout all history path probabilities, such that the summation in **Supplementary Equation (2)** does not require the alignment of the scaling factors before addition. We need to take care when comparing path probabilities though, but this way, we keep the computations precise with almost no overhead in computation time.

#### 2.3 Imputation Core Improvements

The imputation core algorithm computes the *set-maximal matches* of a target haplotype and the reference, for which purpose the reference is reduced to common sites only. As for the phasing part, we keep the basic concept of the original PBWT tool in *EagleImp*, but introduce significant changes in the computation of the set-maximal matches, the handling of missing data, dosage calculation and imputation information (such as allele frequencies and imputation score).

##### 2.3.1 Computation of set-maximal matches

A *match* is defined by an interval $\left[m^{\text{start}}, m^{\text{end}}\right)$ (inclusive start and exclusive end positions of the match in the reduced panel) where the target haplotype matches a set of reference haplotypes exactly. Due to the use of the PBWT data structure, the set can be stored as an interval in the PBWT. We recall the original definition by Durbin [2] here, that a match is set-maximal to a sequence in the reference, if the match is locally maximal, i.e. there is no longer match including this one, and there is also no other sequence in the reference with a longer match that includes this region.

In order to find the set-maximal matches, we create a binary tree data structure containing PBWT intervals in each node. Briefly explained, the algorithm sweeps over all sites and updates the tree by extending all subsequences represented by the PBWT intervals with 0 or 1 according to the current target allele. In particular, all nodes in the tree are processed recursively with each new position, i.e. the interval is mapped as described in **Section 2.1.2**. The complete interval $[0, N-1]$ is inserted as root and the old root is attached as a child. This way, the tree is extended and each node represents an interval that is a subinterval of its parent.

At the point where an interval may not be further extended, the subsequence represented by it is reported as a match and the node is deleted. If the match is reported from the deepest node in the tree it is set-maximal because all other reported matches from this step must be shorter due to later start points and all nodes still alive do not represent a subsequence of this match anymore as they have a later endpoint now. (For further information on the tree structure see detailed explanation below.)

To save computation time, nodes representing equal intervals are merged such that the same interval does not need to be mapped multiple times. However, the nodes keep track of their depth level before merging.

It is clearly to see that all nodes have to be linked in a chain if the target alleles are well defined. The tree represents a simple list in that case. However, the tree representation has its justification as missing data would violate above condition. With a missing allele both extensions are allowed at that site and generate two disjunctive intervals (see below). Also, we leave the opportunity to allow errors in the set-maximal matches. (This option is implemented, but not activated per default as first benchmarks did not show a quality improvement for the extra runtime costs. We leave this open for further investigation.)

**Tree structure for calculating set-maximal matches** For calculating set-maximal matches a PBWT structure similar to that for the condensed reference is created preliminary. In contrast to the PBWTs for the condensed references for each target, this PBWT is slightly larger (because it targets the complete reference without reduction to the $K$ best haplotypes and it misses the reduction to call sites and segments in between). But as the same PBWT can be reused for each target haplotype, the computation is not performance critical.

We use our idea of interval mapping from the phasing part and create a binary tree structure that is updated recursively (depth-first) in every step while iterating $m$ from the first to the last position. A tree node mainly consists of a matching interval $I$ and potentially two subtrees where the root nodes (and therefor all other descending nodes) contain intervals that are a subset of $I$. For convenience, we also add the current depth of the node in the tree, which directly reflects the length of the match represented by the interval. When switching the position from $m$ to $m+1$, we insert a new node with the complete interval $[0, N-1]$ into the tree as root node and the existing tree is attached w.l.o.g. as left tree. With a depth-first strategy all intervals in the nodes are mapped in the PBWT according to the strategy described in the main paper. Mapping always creates two intervals (the 0-interval and the 1-interval, depending on the allele of the corresponding reference sequences at $m+1$). The current interval of the node is updated to the mapping that fits the current haplotype allele which is being matched. Furthermore, the node is attached to its root as a left node, if the selected interval is the 0-interval, and attached to its root as a right node otherwise. A special case occurs, if the current haplotype allele is missing. In this case, a new node is created and both intervals are attached to the root. (Note that the original *PBWT* tool [4] treats missing alleles as reference alleles (0) per default.) At this point we could allow a certain number of mismatches in a set-maximal match by tracking intervals that introduce an error (and not extending the maximal number of allowed errors). However, we did not follow this strategy any further since first tests showed that for allowing a single error the runtime already increased significantly due to the larger tree size, and the imputation quality could not be increased likewise.

A tree node is deleted whenever the corresponding interval cannot be mapped according to the matching haplotype. Descendant subtrees are deleted all the same as if an interval cannot be mapped then clearly its subintervals can neither be. However, at this point, the algorithm has found a potential set-maximal-match at the deepest node of the subtree to be deleted. We used "potential" here since during depth-first processing it is not clear at this point, that the deleted subtree reached the deepest point in the complete tree.

After the recursion returned to the initial caller, all reported matches end at position $m$. Only the longest match is a final candidate for a set-maximal match and needs to be compared to the final depth of the tree. Only if the length is greater or equal to the tree depth, it can be ensured that no match endures in the tree that begins at a position less or equal to that of the match, which makes it set-maximal at last. In order to save computation time, we attached the current tree depth (at the time of starting the recursion) as parameter to the recursion procedure. When it comes to reporting a potential set-maximal match candidate, the length is compared to this parameter before reporting it. This way, we can ensure that we report only the match from the deepest node in the tree which makes a comparison of all match candidates after the recursion obsolete. We introduced another runtime improvement by merging nodes that have a single descendant with the same interval (with tracking the depth of the deeper node). This prevents equal intervals to be mapped multiple times and reduces descents in the tree.

The runtime complexity of this algorithm is $\mathcal{O}(d_m)$ for each position $m$ whereby $d_m$ is the number of nodes in the tree which reflects the number of matching intervals at position $m$. (In most cases, without the presence of missing haplotypes or allowed match errors, this equals the distance of the longest match until here which is the same as the depth of the tree.)

##### 2.3.2 Missing target data

The default reference imputation mode of PBWT handles missing target haplotypes by simply setting these to the reference allele. By explicitly adding the option to impute missing alleles (`--imputeMissing`), PBWT performs a preliminary step before reference imputation and imputes the missing variants using data from the phased target only instead of data from the reference panel.

In *EagleImp*, we simply exclude missing positions from our procedure to compute set-maximal matches, such that these positions are imputed as all other missing variants. In particular, our tree-based procedure described above allows both extensions (0 and 1) at missing sites, which effectively neutralises this position by allowing both alleles.

##### 2.3.3 Calculation of Allele Dosage, Genotype Dosage and Genotype Probabilities

For each variant allele that needs to be imputed, the imputation dosage is calculated as follows. For each sequence (indexed with $i$) from a set-maximal match that covers this variant, a score $s_i$ is computed that depends on the position of the variant in the match and the length of the match:

$$s_i = \left(m - m_i^{\text{start}} + 1\right)\left(m_i^{\text{end}} - m\right) \tag{10}$$

whereby $m$ is the position of the last common variant in the reduced reference panel and $m_i^{\text{start}}$ and $m_i^{\text{end}}$ the start and end positions of the corresponding set-maximal match. Note, that, in contrast to original *PBWT*, we add 1 in the first factor, such that all variants in the first segment in a set-maximal match (between the first and the second common site in the reduced panel) generate a positive score instead of zero.

Then, the allele dosage is simply calculated as:

$$d = \frac{\sum\{s_i | i \text{ with allele at } p \text{ is alt.}\}}{\sum s_i} \tag{11}$$

Exceptions are for variants that were genotyped prior to imputation (which are tagged with TYPED in the output). In this case, allele dosages are 0.0 or 1.0 for homozygous calls. For heterozygous calls, the allele dosage corresponds to the phasing confidence. In the case that variants are not covered by set-maximal matches, the dosage is set to the allele frequency in the reference panel (which is the same approach as in the original PBWT tool).

The imputed allele is selected according to the calculated allele dosage: 0 if $d \leq 0.5$, 1 otherwise.

Furthermore, the user is able to select which fields in the VCF output should accompany the imputed alleles: allele dosages, genotype dosage, genotype probabilities, a combination of those or none. This may significantly reduce the size of the output files if fields that are not necessarily required are omitted. The default is to provide only allele dosages with the genotypes as these form the base for the other values. The genotype dosage is simply the sum of the maternal and paternal allele dosages:

$$gd = d_{\texttt{mat}} + d_{\texttt{pat}} \tag{12}$$

The genotype probabilities are calculated as follows:

$$gp_{\text{homref}} = (1 - d_{\texttt{mat}})(1 - d_{\texttt{pat}}) \tag{13}$$

$$gp_{\text{homalt}} = d_{\texttt{mat}} d_{\texttt{pat}} \tag{14}$$

$$gp_{\text{het}} = 1 - (gp_{\text{homref}} + gp_{\text{homalt}}) \tag{15}$$

##### 2.3.4 Imputation information

In contrast to original *PBWT*, we compute the estimated allele frequency (AF) and the minor allele frequency (MAF) in addition to the allele count (AC) and the allele number (AN) for each variant and provide this information in our output VCF/BCF-file. We also replace the imputation *INFO* score by the imputation $r^2$ calculated as in *minimac4* [3].

$$\text{AF} = \frac{\sum d_n}{N} \tag{16}$$

$$\text{MAF} = \begin{cases} \text{AF} & \text{if AF } \leq 0.5 \\ 1 - \text{AF} & \text{else} \end{cases} \tag{17}$$

$$r^2 = \frac{\sum d_n^2 - \sum d_n \sum d_n}{\sum d_n \left(N - \sum d_n\right)} \tag{18}$$

Here, $d_n$ denote the allele dosages for the current variant and $N$ is the total number of alleles in this dataset. Note, if the divisor in **Supplementary Equation (18)** is zero, $r^2$ is also set to zero.

##### 2.3.5 Multi-processing in imputation

As the imputation of a target sample is independent from other targets, a simple parallelisation strategy is to run multiple threads for imputing several targets concurrently. We perform this strategy for the computation of set-maximal matches. However, when it comes to the actual imputation the variant-major format of the output files presents a major problem as for each variant the imputed data of all targets is required at once and writing the output via the *HTSlib* is slow. We address this problem with the following approach.

First of all, we launch a fourth of the available system threads as *writer threads* with each writer creating a temporary output file (i.e. 8 threads when 32 threads are available). All output variants are evenly distributed in the same number of coherent blocks with each block assigned to one output file. Each block is then evenly distributed in coherent chunks with a variable number of variants that is derived from the number of samples. (In particular, the chunk size times the number of samples is 10 million. This value delivered good results but may be subject for optimisation.)

Each chunk is processed using the remaining threads to impute its variants parallelised over the targets. The blocks and chunks are processed alternating over the blocks (i.e. first chunk from first block, then first chunk from second block etc.). Each chunk directs the imputation output to the writer queue of the corresponding output thread assigned to the current block. This way, the load to the output queues is balanced and the multi-processing capabilities of the computing machine can be fully used.

After the imputation the generated temporary output files are simply concatenated to a single file. As the packed *VCF* or *BCF* files are in *Block Gzip Format (BGZF)* format, this is possible without any difficulty as packing is handled block-wise (but would have not been possible if the output format was standard *zip*). **Figure 2** in the main paper illustrates the multi-processing scheme for imputation.

### 3 Benchmarks

#### 3.1 Preparation of three reference panels

First benchmarks were performed using the Haplotype Reference Consortium (HRC) Release 1.1 [6] portion of the SIS imputation reference (hereafter referred to as the *HRC1.1*), which is available through the European Genome-phenome Archive (EGA) upon user request (approved and downloaded on Oct 26, 2020). This part of the HRC panel contains 40.4 million variants for chromosomes 1 to 22 and chromosome X, with the number of samples being 22,691 for chromosome 1, 26,199 for chromosome X (including 11,739 males), and 27,165 for all other chromosomes. The second benchmark panel was the 1000 Genomes (1000G) Phase 3 reference panel [7] with the latest updates to version 5a from Jul 31, 2020 (hereafter referred to as the *1000G Phase 3*). This panel contains 84.8 million variants for chromosomes 1 to 22 and chromosomes X and Y, for a total of 2,504 samples (including 1,233 male samples). To demonstrate that *EagleImp* can use large reference panels in the order of one million samples for phasing and imputation in the future, we artificially synthesised a third benchmark panel (hereafter referred to as the *synthetic HRC1.1*) based on the *HRC1.1* panel described above. We used *bcftools* [8] to increase the number of samples in the panel by a factor of 40 (repeated use of `bcftools merge --force-samples`). The result was a panel with more than one million samples (>2 million haplotypes), in particular 907,640 samples for chromosome 1, 1,047,960 samples for chromosome X and 1,086,600 samples for all other chromosomes.

#### 3.2 Preparation of 18 target benchmark datasets

For our benchmarks with the *HRC1.1* panel, we created six sub-datasets as target files (i.e. GWAS input files) from *HRC1.1*: First, we selected all 494 samples of European ancestry (*HRC.EUR*) that were also included in the *1000G Phase 3* panel. Next, we selected another five target sets of 500 randomly selected samples each (*HRC.v1* to *HRC.v5*). Since chromosome 1 contained fewest samples, we selected the samples from the chromosome 1 dataset. To create a realistic scenario for phasing and imputation, we then reduced the genomic information of the target datasets to the genetic variants contained on Illumina's *Global Screening Array (GSA)* [9], which combines multi-ethnic genome-wide content, curated clinical research variants, and quality control (QC) markers for precision medicine research; further, we removed the phase information to create unphased genotypes. (In fact, the phase information did not have to be explicitly removed as both *EagleImp* and *Eagle2* ignore this information by default when reading the target input file for the phasing process.) The final benchmark target datasets contained 619,872 variants found on both the GSA and the *HRC1.1* panel. For each of the six target datasets, we further reduced the *HRC1.1* panel by the samples from the respective target datasets to avoid duplicate samples in target and reference panel.

Similary, using the *1000G Phase 3* panel, we created ten further benchmark datasets. For five of the ten benchmark sets, we randomly selected 50 samples of European ancestry (*1kG.EUR.v1* to *1kG.EUR.v5*), and for the other five, we randomly selected 50 samples from the whole panel (*1kG.v1* to *1kG.v5*). As for the *HRC1.1* panel benchmark sets, we reduced the set of variants to the GSA marker content (resulting in 647,963 variants shared between the GSA and the *1000G Phase 3* panel) and created a reduced *1000G Phase 3* panel without the target samples for each benchmark dataset.

To compare the imputation accuracy of *EagleImp* with the accuracy of current imputation servers (SIS, MIS and Topmed), we chose two real-world target GWAS datasets from GSA genotyping named *COVID.Italy* and *COVID.Spain* from a COVID-19 Genome-wide Association Study (GWAS) [10] comprising 2,113 samples (839 cases, 1,274 controls) typed at 559,519 variants and 1,792 samples (842 cases, 950 controls) typed at 549,696 variants, respectively, and performed phasing and imputation with our *HRC1.1* reference panel. Finally, we used the larger set, *COVID.Italy*, also to demonstrate the capabilities of *EagleImp* with large reference panels by imputing it against our *synthetic HRC1.1* panel with >1 million samples.

#### 3.3 Benchmark parameters

Quality benchmarks were performed for six datasets with the HRC1.1 reference panel subset (*HRC.EUR*, *HRC.v1-5*), and for ten datasets for the 1000 Genomes reference panel (*1kG.v1-5*, *1kG.v1-5.EUR*). The runtime benchmarks were performed with several multi-processing configurations for *EagleImp* and *Eagle2* on the *HRC.EUR* dataset. For *PBWT* we simply processed all chromosome datasets in parallel as *PBWT* lacks support for multi-threading.

The genetic map used for phasing in *EagleImp* and *Eagle2* was downloaded from: `https://alkesgroup.broadinstitute.org/Eagle/downloads/tables/`

We used the following runtime parameters for *Eagle2*:

```
eagle --geneticMapFile <genetic map>
      --outPrefix <output file prefix>
      --vcfOutFormat b --vcfRef <reference bcf file>
      --vcfTarget <target input bcf file>
      --allowRefAltSwap --noImpMissing
      --numThreads <number of threads>
      --Kpbwt <K> --pbwtIters 1
```

And we used the following runtime parameters for *PBWT*:

```
pbwt -readVcfGT <phased output bcf file>
     -referenceImpute <reference pbwt file>
     -writeBcfGz <output bcf file>
```

The following parameters were activated for *EagleImp*:

```
eagleimp --geneticMap <genetic map>
         -o <output file prefix>
         --vcfOutFormat b --ref <reference qref file>
         --target <target input bcf file>
         --imputeInfo 'a' --outputPhasedFile
         --stat <status file>
         --allowRefAltSwap
         -t <number of threads>
         -K <K> -i 1
```

#### 3.4 The impact of pre-phasing and reverse phasing

The following **Supplementary Tables 1**, **2** and **3** show the impact of pre-phasing and reverse phasing exemplarily for the *HRC.** datasets and $K = 10,000$ and $K = \max$ for *EagleImp*. The results are illustrated in **Supplementary Figure 2**. Note, that we disabled pre-phasing for our following benchmarks as it generates slightly better results.

Supplementary Table 1: Phasing switch error rates of *HRC.EUR* dataset for *EagleImp* using different combinations of pre-phasing and reverse phasing disabled/enabled.

| | $K = 10,000$ | $K = 16,384$ | $K = 32,768$ | $K = \max$ |
|---|---|---|---|---|
| no pre-phasing | 0.0047019 | 0.0045232 | 0.0043352 | 0.0042354 |
| pre-ph. + rev.ph. | 0.0047052 | 0.0045258 | 0.0043365 | 0.0042366 |
| no reverse phasing | 0.0048520 | 0.0046750 | 0.0044891 | 0.0043929 |

Supplementary Table 2: Imputation genotype error rates of *HRC.EUR* dataset for *EagleImp* using different combinations of pre-phasing and reverse phasing disabled/enabled.

| | $K = 10,000$ | $K = 16,384$ | $K = 32,768$ | $K = \max$ |
|---|---|---|---|---|
| no pre-phasing | 0.0026500 | 0.0025999 | 0.0025415 | 0.0025021 |
| pre-ph. + rev.ph. | 0.0026517 | 0.0026012 | 0.0025439 | 0.0025036 |
| no reverse phasing | 0.0026770 | 0.0026260 | 0.0025671 | 0.0025274 |

Supplementary Table 3: Runtimes (in seconds) of *HRC.EUR* dataset in *2x4x8* multi-processing configuration for *EagleImp* using different combinations of pre-phasing and reverse phasing disabled/enabled.

| | $K = 10,000$ | $K = 16,384$ | $K = 32,768$ | $K = \max$ |
|---|---|---|---|---|
| no pre-phasing | 1299 | 1507 | 2061 | 2125 |
| pre-ph. + rev.ph. | 1329 | 1505 | 2112 | 2102 |
| no reverse phasing | 1199 | 1388 | 2003 | 1969 |

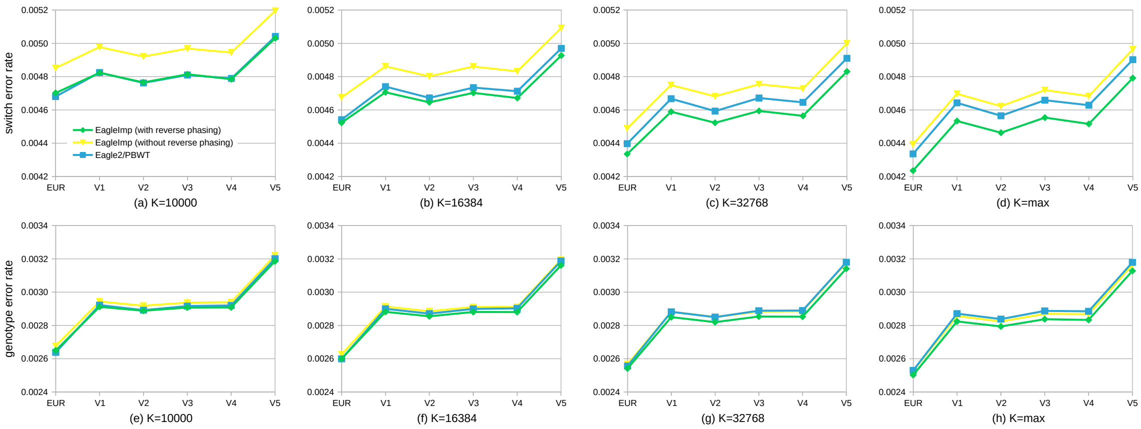

Supplementary Figure 2: Phasing switch error rates and imputation genotype error rates of the *HRC.EUR* and *HRC.v1–5* datasets with reverse phasing enabled (green) and not enabled (yellow) in comparison to original Eagle2/PBWT (blue).

##### 3.5 Switch errors and genotype errors

Supplementary Table 4: Phasing switch error rates of *HRC.** datasets.

| | | HRC.EUR | HRC.v1 | HRC.v2 | HRC.v3 | HRC.v4 | HRC.v5 |
|---|---|---|---|---|---|---|---|
| $K = 10,000$ | Eagle2 | 0.0046801 | 0.0048247 | 0.0047625 | 0.0048100 | 0.0047895 | 0.0050417 |
| | EagleImp | 0.0047019 | 0.0048231 | 0.0047655 | 0.0048144 | 0.0047849 | 0.0050312 |
| $K = 16,384$ | Eagle2 | 0.0045413 | 0.0047402 | 0.0046724 | 0.0047342 | 0.0047123 | 0.0049699 |
| | EagleImp | 0.0045232 | 0.0047059 | 0.0046457 | 0.0047026 | 0.0046715 | 0.0049273 |
| $K = 32,768$ | Eagle2 | 0.0043971 | 0.0046674 | 0.0045930 | 0.0046716 | 0.0046456 | 0.0049103 |
| | EagleImp | 0.0043352 | 0.0045892 | 0.0045228 | 0.0045939 | 0.0045644 | 0.0048310 |
| $K = \max$ | Eagle2 | 0.0043361 | 0.0046427 | 0.0045648 | 0.0046583 | 0.0046284 | 0.0049021 |
| | EagleImp | 0.0042354 | 0.0045334 | 0.0044631 | 0.0045540 | 0.0045160 | 0.0047917 |

Supplementary Table 5: Imputation genotype error rates of *HRC.** datasets.

| | | HRC.EUR | HRC.v1 | HRC.v2 | HRC.v3 | HRC.v4 | HRC.v5 |
|---|---|---|---|---|---|---|---|
| $K = 10,000$ | Eagle2/PBWT | 0.0026380 | 0.0029222 | 0.0028917 | 0.0029163 | 0.0029198 | 0.0032014 |
| | EagleImp | 0.0026500 | 0.0029118 | 0.0028882 | 0.0029075 | 0.0029083 | 0.0031857 |
| $K = 16,384$ | Eagle2/PBWT | 0.0025994 | 0.0028999 | 0.0028708 | 0.0028993 | 0.0029028 | 0.0031873 |
| | EagleImp | 0.0025999 | 0.0028815 | 0.0028549 | 0.0028807 | 0.0028802 | 0.0031611 |
| $K = 32,768$ | Eagle2/PBWT | 0.0025563 | 0.0028818 | 0.0028499 | 0.0028884 | 0.0028894 | 0.0031797 |
| | EagleImp | 0.0025415 | 0.0028500 | 0.0028198 | 0.0028535 | 0.0028525 | 0.0031412 |
| $K = \max$ | Eagle2/PBWT | 0.0025298 | 0.0028708 | 0.0028381 | 0.0028874 | 0.0028850 | 0.0031790 |
| | EagleImp | 0.0025021 | 0.0028240 | 0.0027938 | 0.0028375 | 0.0028331 | 0.0031278 |

Supplementary Table 6: Phasing switch error rates of *1kG.** datasets.

| | 1kG.v1 | 1kG.v2 | 1kG.v3 | 1kG.v4 | 1kG.v5 |
|---|---|---|---|---|---|
| Eagle2 | 0.0069949 | 0.0068818 | 0.0073099 | 0.0070385 | 0.0071055 |
| EagleImp | 0.0067150 | 0.0066103 | 0.0069323 | 0.0067644 | 0.0067597 |

Supplementary Table 7: Phasing switch error rates of *1kG.EUR.** datasets.

| | 1kG.EUR.v1 | 1kG.EUR.v2 | 1kG.EUR.v3 | 1kG.EUR.v4 | 1kG.EUR.v5 |
|---|---|---|---|---|---|
| Eagle2 | 0.0054347 | 0.0059278 | 0.0057152 | 0.0056280 | 0.0054168 |
| EagleImp | 0.0052382 | 0.0057262 | 0.0055194 | 0.0054426 | 0.0052236 |

Supplementary Table 8: Imputation genotype error rates of *1kG.** datasets.

| | 1kG.v1 | 1kG.v2 | 1kG.v3 | 1kG.v4 | 1kG.v5 |
|---|---|---|---|---|---|
| Eagle2/PBWT | 0.0055262 | 0.0053962 | 0.0057381 | 0.0052965 | 0.0055669 |
| EagleImp | 0.0054702 | 0.0053468 | 0.0056495 | 0.0052496 | 0.0055025 |

Supplementary Table 9: Imputation genotype error rates of *1kG.EUR.** datasets.

| | 1kG.EUR.v1 | 1kG.EUR.v2 | 1kG.EUR.v3 | 1kG.EUR.v4 | 1kG.EUR.v5 |
|---|---|---|---|---|---|
| Eagle2/PBWT | 0.0036921 | 0.0038560 | 0.0037806 | 0.0037664 | 0.0036922 |
| EagleImp | 0.0036640 | 0.0038275 | 0.0037587 | 0.0037385 | 0.0036665 |

##### 3.6 Imputation Accuracy $r^2$ in COVID-19 GWAS Data

For the real-world GWAS datasets *COVID.Italy* and *COVID.Spain* (2,113 Italian and 1,792 Spanish samples (severe COVID-19 cases plus controls) genotyped at around 550,000 variants), we computed imputation $r^2$ values stratified by MAF (including all variants above this threshold) using the four different $K$ parameters from above. First, we extracted $r^2$ values and estimated MAFs of imputed variants from the *EagleImp* results. For the *Eagle2/PBWT* results, we needed to calculate imputation MAF and $r^2$ from reported allele dosages (**Supplementary Equation (18)**), as these are not included in the *PBWT* output. Since there is no sample overlap between our real-world GWAS datasets and the *HRC1.1* reference panel provided by the imputation servers *MIS* [11] and *SIS* [12] (32,470 samples of predominantly European ancestry; this larger HRC1.1 panel from *MIS* and *SIS* is not freely available [6]), we further phased and imputed both GWAS datasets once with the *MIS* and once with the *SIS* and the full HRC panel to include these results in the comparison. In addition, we imputed both GWAS datasets using the *TOPMed* imputation server [13] with its customised TOPMed panel [14], which consists of 97,256 reference samples (release 2) compiled from more than 85 WGS studies. Thus, the *TOPMed* imputation accuracy (as a reference model) shows us the current limits of imputation quality (independent of the underlying algorithm), which is due to the enormous size of the reference panel. For *MIS* and *TOPMed*, we extracted MAF and $r^2$ directly from imputation results, as we did for *EagleImp*. Since the *SIS* uses original *PBWT* for imputation, we had to recalculate $r^2$ values as described above.

The resulting $r^2$-plots are depicted in the following **Supplementary Figures 3** and **4**.

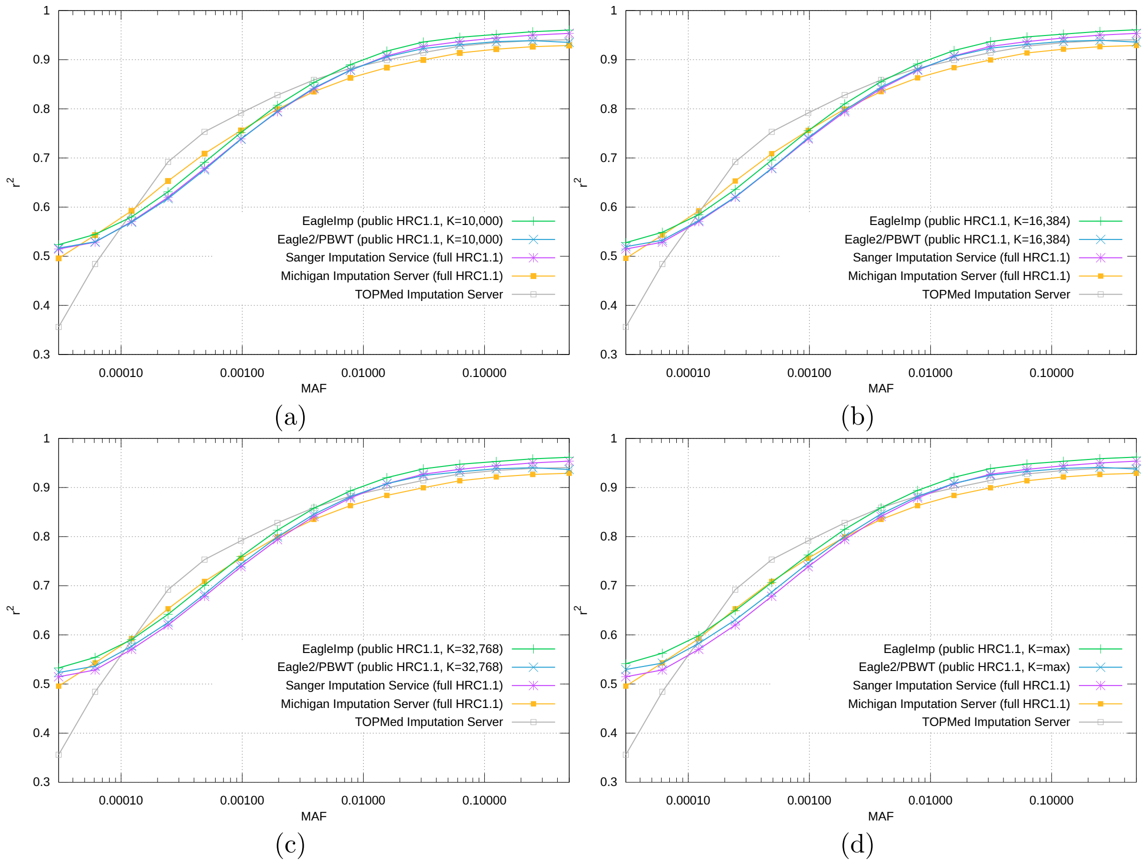

Supplementary Figure 3: $r^2$-plots for the *COV.Italy* dataset using different values of $K$ for phasing. (a) $K = 10,000$, (b) $K = 16,384$, (c) $K = 32,768$, (d) $K = \max$

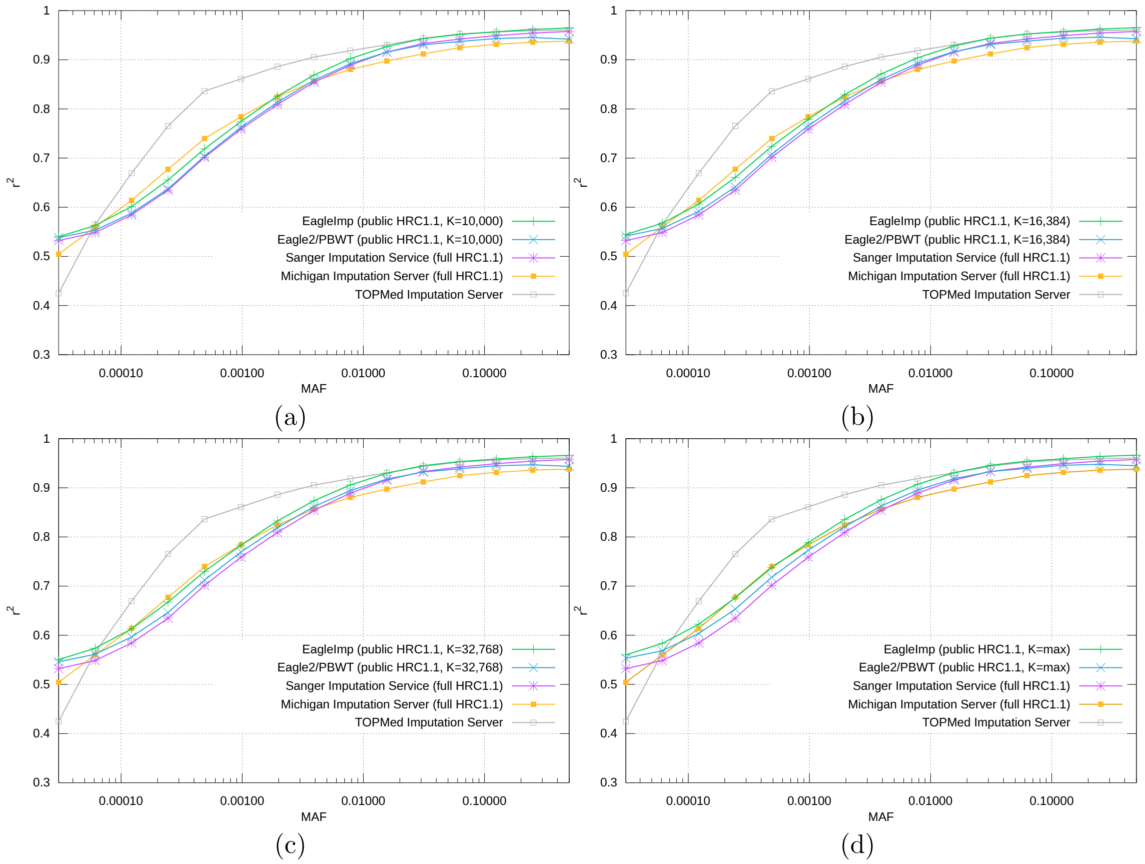

Supplementary Figure 4: $r^2$-plots for the $COV.Spain$ dataset using different values of $K$ for phasing. (a) $K = 10,000$, (b) $K = 16,384$, (c) $K = 32,768$, (d) $K = \max$

#### 3.7 Multi-processing Runtimes

The different multi-processing runtimes are distinguished by the number of concurrent worker processes started and the number of threads assigned to each worker. An *Eagle2/PBWT* run in *8x4* configuration means that 8 concurrent worker processes are executed in parallel with 4 threads assigned to each worker.

For *EagleImp* the number of worker processes that share a CPU-lock is also relevant. A *2x4x8* configuration means that 2x4 workers are executed in parallel with each two workers sharing a CPU-lock, and 8 threads (mutual exclusion via the lock) are assigned to each worker.

Supplementary Table 10: Runtimes (in seconds) for the complete *HRC.EUR* dataset for different multi-processing configurations.

| | | $K = 10,000$ | $K = 16,384$ | $K = 32,768$ | $K = \max$ |
|---|---|---|---|---|---|
| Eagle2/PBWT | 1x32 | 6386 | 6741 | 7733 | 8830 |
| | 2x16 | 4110 | 4382 | 5240 | 6322 |
| | 4x8 | 3068 | 3464 | 4352 | 5388 |
| | 8x4 | 2778 | 3148 | 4131 | 5131 |
| | 16x2 | 2783 | 3184 | 4240 | 5419 |
| | 32x1 | 3257 | 3870 | 5590 | 7122 |
| EagleImp | 1x32 | 2401 | 2627 | 3203 | 3260 |
| | 2x1x32 | 1933 | 2081 | 2681 | 2767 |
| | 2x16 | 1611 | 1816 | 2351 | 2363 |
| | 2x2x16 | 1401 | 1630 | 2192 | 2192 |
| | 4x8 | 1354 | 1579 | 2154 | 2126 |
| | 2x4x8 | 1299 | 1507 | 2061 | 2125 |
| | 8x4 | 1329 | 1521 | 2092 | 2064 |
| | 2x8x4 | 1358 | 2028 | 2212 | 2387 |

#### 3.8 Chromosome-wise Runtimes of Separated Steps

Supplementary Table 11: Phasing only runtimes (in seconds) for the single chromosomes of the *HRC.EUR* dataset in *1x32* configuration and different values of $K$.

| | Eagle2/PBWT | | | | EagleImp | | | |
|---|---|---|---|---|---|---|---|---|
| | $K = 10,000$ | $K = 16,384$ | $K = 32,768$ | $K = \max$ | $K = 10,000$ | $K = 16,384$ | $K = 32,768$ | $K = \max$ |
| chr1 | 346 | 372 | 442 | 493 | 83 | 100 | 145 | 134 |
| chr2 | 424 | 453 | 536 | 616 | 91 | 110 | 157 | 160 |
| chr3 | 356 | 381 | 442 | 517 | 76 | 90 | 130 | 133 |
| chr4 | 345 | 366 | 425 | 497 | 72 | 86 | 124 | 126 |
| chr5 | 320 | 344 | 399 | 455 | 69 | 80 | 116 | 119 |
| chr6 | 328 | 349 | 421 | 499 | 73 | 89 | 130 | 133 |
| chr7 | 285 | 307 | 362 | 423 | 63 | 74 | 107 | 112 |
| chr8 | 279 | 294 | 345 | 402 | 58 | 68 | 99 | 103 |
| chr9 | 217 | 232 | 273 | 325 | 50 | 59 | 86 | 88 |
| chr10 | 249 | 266 | 313 | 370 | 56 | 67 | 96 | 99 |
| chr11 | 252 | 264 | 314 | 366 | 54 | 64 | 93 | 98 |
| chr12 | 239 | 254 | 302 | 354 | 54 | 64 | 93 | 97 |
| chr13 | 182 | 191 | 226 | 270 | 41 | 48 | 70 | 73 |
| chr14 | 163 | 176 | 202 | 243 | 37 | 44 | 62 | 65 |
| chr15 | 154 | 161 | 194 | 225 | 37 | 44 | 61 | 64 |
| chr16 | 166 | 177 | 211 | 248 | 42 | 49 | 67 | 71 |
| chr17 | 145 | 155 | 184 | 218 | 36 | 43 | 60 | 63 |
| chr18 | 144 | 154 | 182 | 214 | 35 | 41 | 58 | 61 |
| chr19 | 116 | 123 | 144 | 173 | 29 | 34 | 47 | 50 |
| chr20 | 119 | 126 | 151 | 183 | 30 | 36 | 50 | 53 |
| chr21 | 73 | 76 | 89 | 107 | 17 | 21 | 29 | 30 |
| chr22 | 72 | 76 | 90 | 106 | 18 | 22 | 31 | 31 |
| chrX | 201 | 209 | 240 | 275 | 35 | 42 | 60 | 68 |

Supplementary Table 12: Imputation only runtimes (in seconds) for the single chromosomes of the *HRC.EUR* dataset in *1x32* configuration and different values of $K$.

| | Eagle2/PBWT | | | | EagleImp | | | |
|---|---|---|---|---|---|---|---|---|
| | $K = 10,000$ | $K = 16,384$ | $K = 32,768$ | $K = \max$ | $K = 10,000$ | $K = 16,384$ | $K = 32,768$ | $K = \max$ |
| chr1 | 964 | 983 | 1093 | 1100 | 115 | 113 | 102 | 115 |
| chr2 | 1194 | 1215 | 1224 | 1232 | 134 | 140 | 138 | 134 |
| chr3 | 1135 | 1161 | 1015 | 1156 | 115 | 116 | 109 | 116 |
| chr4 | 1095 | 1100 | 959 | 1061 | 114 | 114 | 111 | 111 |
| chr5 | 1025 | 1012 | 999 | 1043 | 103 | 103 | 103 | 103 |
| chr6 | 1010 | 926 | 954 | 974 | 108 | 107 | 101 | 108 |
| chr7 | 920 | 881 | 825 | 913 | 95 | 94 | 95 | 88 |
| chr8 | 934 | 830 | 832 | 827 | 90 | 89 | 89 | 84 |
| chr9 | 685 | 666 | 642 | 691 | 71 | 70 | 69 | 71 |
| chr10 | 745 | 794 | 761 | 839 | 78 | 79 | 80 | 78 |
| chr11 | 703 | 737 | 850 | 810 | 78 | 79 | 80 | 76 |
| chr12 | 819 | 798 | 709 | 725 | 77 | 77 | 78 | 77 |
| chr13 | 601 | 570 | 638 | 584 | 59 | 57 | 56 | 58 |
| chr14 | 648 | 611 | 625 | 460 | 53 | 53 | 53 | 52 |
| chr15 | 444 | 479 | 511 | 460 | 50 | 50 | 50 | 47 |
| chr16 | 440 | 588 | 638 | 442 | 56 | 56 | 55 | 55 |
| chr17 | 407 | 370 | 444 | 418 | 46 | 46 | 47 | 47 |
| chr18 | 497 | 515 | 461 | 405 | 46 | 46 | 46 | 45 |
| chr19 | 309 | 381 | 384 | 491 | 38 | 38 | 38 | 37 |
| chr20 | 446 | 373 | 387 | 318 | 38 | 39 | 39 | 38 |
| chr21 | 291 | 243 | 182 | 206 | 22 | 23 | 23 | 23 |
| chr22 | 304 | 182 | 210 | 194 | 24 | 23 | 23 | 23 |
| chrX | 518 | 433 | 446 | 509 | 52 | 53 | 53 | 53 |

Supplementary Table 13: Combined (phasing and imputation) runtimes (in seconds) for the single chromosomes of the *HRC.EUR* dataset in *1x32* configuration and different values of $K$.

| | Eagle2/PBWT | | | | EagleImp | | | |
|---|---|---|---|---|---|---|---|---|
| | $K = 10,000$ | $K = 16,384$ | $K = 32,768$ | $K = \max$ | $K = 10,000$ | $K = 16,384$ | $K = 32,768$ | $K = \max$ |
| chr1 | 1310 | 1355 | 1535 | 1592 | 167 | 182 | 217 | 218 |
| chr2 | 1618 | 1668 | 1760 | 1848 | 185 | 209 | 255 | 255 |
| chr3 | 1491 | 1541 | 1456 | 1673 | 159 | 173 | 207 | 217 |
| chr4 | 1439 | 1466 | 1383 | 1558 | 154 | 168 | 203 | 207 |
| chr5 | 1344 | 1356 | 1398 | 1497 | 143 | 155 | 189 | 194 |
| chr6 | 1338 | 1275 | 1375 | 1473 | 150 | 165 | 200 | 211 |
| chr7 | 1205 | 1187 | 1187 | 1336 | 131 | 141 | 176 | 174 |
| chr8 | 1212 | 1124 | 1177 | 1229 | 123 | 133 | 164 | 161 |
| chr9 | 902 | 898 | 915 | 1016 | 101 | 109 | 134 | 140 |
| chr10 | 994 | 1061 | 1074 | 1209 | 111 | 123 | 153 | 154 |
| chr11 | 955 | 1001 | 1165 | 1175 | 109 | 121 | 151 | 151 |
| chr12 | 1057 | 1052 | 1011 | 1079 | 109 | 119 | 149 | 152 |
| chr13 | 783 | 761 | 864 | 854 | 83 | 89 | 109 | 114 |
| chr14 | 811 | 787 | 826 | 702 | 75 | 82 | 100 | 101 |
| chr15 | 597 | 641 | 705 | 685 | 73 | 79 | 97 | 97 |
| chr16 | 606 | 765 | 849 | 690 | 79 | 87 | 105 | 109 |
| chr17 | 552 | 525 | 628 | 636 | 69 | 76 | 94 | 97 |
| chr18 | 641 | 669 | 643 | 619 | 68 | 74 | 91 | 93 |
| chr19 | 425 | 504 | 529 | 663 | 57 | 61 | 75 | 76 |
| chr20 | 564 | 500 | 538 | 500 | 57 | 63 | 78 | 81 |
| chr21 | 364 | 319 | 271 | 313 | 34 | 38 | 47 | 46 |
| chr22 | 376 | 258 | 300 | 300 | 36 | 39 | 48 | 48 |
| chrX | 719 | 642 | 686 | 784 | 71 | 78 | 96 | 104 |

#### 3.9 Single-thread Runtimes

Supplementary Table 14: Single-thread runtimes (in seconds) for chromosome 2 from *HRC.EUR* dataset.

| | | $K = 10,000$ | $K = 16,384$ | $K = 32,768$ | $K = \max$ |
|---|---|---|---|---|---|
| Eagle2/PBWT | phasing | 2015 | 2546 | 3918 | 5460 |
| | imputation | 1123 | 1152 | 1187 | 1164 |
| | total | 3138 | 3698 | 5105 | 6624 |
| EagleImp | phasing | 997 | 1373 | 2303 | 2198 |
| | imputation | 800 | 796 | 789 | 792 |
| | combined* | 1753 | 2125 | 3047 | 2946 |

* combined runtime is less than the sum of phasing and imputation as the reference panel does not need to be loaded twice in *EagleImp*.

Supplementary Table 15: Single-thread runtimes (in seconds) for chromosome 21 from *HRC.EUR* dataset.

| | | $K = 10,000$ | $K = 16,384$ | $K = 32,768$ | $K = \max$ |
|---|---|---|---|---|---|
| Eagle2/PBWT | phasing | 373 | 473 | 730 | 1012 |
| | imputation | 186 | 186 | 185 | 185 |
| | total | 559 | 659 | 915 | 1197 |
| EagleImp | phasing | 205 | 273 | 460 | 443 |
| | imputation | 124 | 125 | 127 | 127 |
| | combined* | 322 | 390 | 580 | 563 |

* combined runtime is less than the sum of phasing and imputation as the reference panel does not need to be loaded twice in *EagleImp*.

#### 3.10 Runtimes of COVID-19 GWAS Data

The benchmarks for the real-world datasets *COVID.Italy* and *COVID.Spain* were performed in *EagleImp.2x4x8* and *Eagle2.8x4* configurations respectively. The runtimes are presented in the following **Supplementary Table 16**.

Supplementary Table 16: Runtimes (in seconds) for *COV.Italy* and *COV.Spain* datasets.

| | | $K = 10,000$ | $K = 16,384$ | $K = 32,768$ | $K = \max$ |
|---|---|---|---|---|---|
| *COV.Italy* | Eagle2/PBWT (8x4) | 9,044 | 10,270 | 14,286 | 18,802 |
| | EagleImp (2x4x8) | 5,260 | 6,164 | 8,589 | 8,855 |
| *COV.Spain* | Eagle2/PBWT (8x4) | 7,443 | 8,662 | 12,008 | 15,578 |
| | EagleImp (2x4x8) | 4,317 | 5,258 | 7,074 | 7,247 |

#### 3.11 Preparation of Reference Data

As *Eagle2* handles reference files in `.vcf.gz` or `.bcf` format, *PBWT* requires these files to be converted in `.pbwt` format. *EagleImp* can handle reference files in `.vcf.gz` or `.bcf` format as well, but we recommend a conversion into our `.qref` format for performance reasons, already if the user intends to do more than only one analysis with the same reference. All presented benchmarks for *EagleImp* in this manuscript use `.qref` references.

In **Supplementary Table 17** we present the preparation times for `.pbwt` and `.qref` files for the 1000 Genomes and HRC panel used in our benchmarks. The table also contains the runtime for creating our synthesised panel with 40 times the samples from the public HRC1.1 panel, which we created by a repetitive execution of `bcftools merge --force-samples`. Note, that we ran all conversions for each chromosome concurrently on our system.

For `.pbwt` creation we used the following command:

```
pbwt -readVcfGT <reference vcf/bcf file>
     -writeAll <reference file prefix>
```

The following parameters were activated for *EagleImp* for creating `.qref` files:

```
eagleimp --ref <reference bcf file>
         --makeQref
```

Supplementary Table 17: Runtimes (in seconds) of creating `.pbwt` and `.qref` files for the used reference panels. The conversions of the individual chromosomes were all started in parallel.

| | `.pbwt` | `.qref` |
|---|---|---|
| 1000 Genomes Phase 3 | 880 | 890 |
| HRC 1.1 (public release) | 4,000 | 4,603 |
| synthetic HRC 1.1 (HRC1.1×40) | 76,216 | 41,211 |

#### 3.12 Large Reference Panel

Supplementary Table 18: Runtimes (in seconds) for *COV.Italy* data and the *synthetic HRC1.1* panel with >1 million samples. $K = 10,000$ is the default in *Eagle2*, $K = 54,330$ is the maximum value for a benchmark against the public *HRC1.1* panel, $K = \max$ corresponds to all >2 million haplotypes.

| | | $K = 10,000$ | $K = 54,330$ | $K = \max$ |
|---|---|---|---|---|
| Eagle2/PBWT | 1x32 | 375,368 | — | 1,314,484 * |
| | 8x4 | 244,245 | 253,125 | failed ** |
| EagleImp | 1x32 | 124,304 | — | 375,281 |
| | 2x4x8 | 87,176 | 86,481 | 346,883 |

* partially failed on chromosomes 1–4,6 due to insufficient memory
** failed completely due to insufficient memory

[8] Wellcome Sanger Institute, BCFTools v1.10.2, https://github.com/samtools/bcftools;.

[9] Illumina, Infinium Global Screening Array-24 Kit, https://www.illumina.com/products/by-type/microarray-kits/infinium-global-screening.html;.

[10] Ellinghaus D, Degenhardt F, Bujanda L, Buti M, Albillos A, Invernizzi P, et al. Genomewide association study of severe COVID-19 with respiratory failure. The New England Journal of Medicine 2020 Oct;383(16):1522–1534.

[11] US National Institutes of Health, Michigan Imputation Server, https://imputationserver.sph.umich.edu/;.

[12] Wellcome Sanger Institute, Sanger Imputation Service, https://imputation.sanger.ac.uk/;.

[13] National Heart, Lung, and Blood Institute, BioData Catalyst TOPMed Imputation Server, https://imputation.biodatacatalyst.nhlbi.nih.gov/;.

[14] Taliun D, Harris DN, Kessler MD, Carlson J, Szpiech ZA, Torres R, et al. Sequencing of 53,831 diverse genomes from the NHLBI TOPMed Program. bioRxiv 2019;https://www.biorxiv.org/content/early/2019/03/06/563866.
